# A Modular In-Incubator Microscope for Longitudinal Live Cell Microscopy

**DOI:** 10.64898/2026.01.20.699789

**Authors:** Drew Ehrlich, Yohei Rosen, Santhosh Arul, John Minnick, Seth Nicholson, Kateryna Voitiuk, Spencer Seiler, Anna Toledo, Samira Vera-Choqqueccota, Neil Doherty, Jess Sevetson, Max McGlynn, Kivilcim Doganyigit, Maryam Moarefian, Sri Kurniawan, Mohammed A. Mostajo-Radji, Sofie R. Salama, Ethan Winkler, David Haussler, Mircea Teodorescu

## Abstract

Longitudinal live cell imaging is valuable for characterizing dynamic morphological and phenotypic changes in biological systems. However, conventional approaches rely on manual microscope operation, which is labor-intensive, limits imaging frequency, and disrupts the cellular environment. These constraints reduce scalability, increase experimental variability, and restrict both the duration and temporal resolution of continuous imaging. Although automated imaging platforms partially address these limitations, existing solutions are often constrained by the cost, footprint, and inflexibility of in-incubator microscopes or stage-top incubators.

Here, we present an automated in-incubator epifluorescence microscope designed for long-term operation. The system features a modular architecture with optional multi-fluorescence imaging, automated plate scanning, configurable light sources, and compatibility with multiple plate formats, including integration with fluidic automation devices. By positioning the light sources and control electronics outside the incubator, the platform improves thermal stability and long-term operational reliability. This approach enables continuous, high-frequency imaging over extended durations, providing a source of rich data for quantifying time-dependent tissue phenotypes, morphological remodeling, and transient biological processes.

## INTRODUCTION

### 1 Introduction

Live-cell microscopy enables direct observation of spatiotemporal dynamics in biological systems, allowing researchers to quantify how tissues evolve over time within their native microenvironments.^1,2^. Long-term, high-frequency imaging captures dynamic cellular behaviors such as migration^3^, cell-cell interactions^4^, cell-matrix interactions, as well as cell state phenotypes including cellular morphological changes^5^, mitoses and apoptoses, differentiation^6^ and fluorophore-conjugated protein expression^7^. These data can be used to infer underlying cellular processes and intercellular interactions^8–10^. By capturing repeated measurements from the same cells over time, time-series imaging enables the detection of temporal correlations, causal relationships, and rare or transient events that are difficult to resolve from cross-sectional snapshots alone. As a result, live-cell imaging provides rich data that improve experimental insight into dynamic mechanisms underlying development, disease progression, and therapeutic response.

In research settings, live cell microscopy equipment is generally limited to three formats: bench microscopes with stage top incubation chambers^11–14^, specialized microscopes that operate inside of cell culture incubators^15,16^, and cell plate ‘hotel’ systems that use a robotic manipulator to remove dishes from an incubation system for imaging^17–19^, automating the standard process by which a technician would take plates out of an incubator for imaging.

Stagetop incubation systems are small incubation chambers that are combined with conventional laboratory microscopes. These can be a valuable tool to observe live samples for short periods^11,20,21^. However, they lack the insulation and sealing of standard free-standing cell culture incubators, resulting in the physiological stresses of evaporative osmotic toxicity, condensation and temperature instability and are also generally size-limited, placing limitations on the form factor of the cell culture vessels inside.

In-incubator microscopes are an alternative to stagetop incubation systems. These microscopes are specifically designed to be used inside a cell culture incubator^15,16,22,23^. To accomplish this, these devices have to be designed to mitigate the thermal loads imposed by the humid and mildly acidic conditions of the cell culture incubator, which can cause both corrosion and microbial growth. Incubator microscopes exist because placing a standard benchtop microscope inside an incubator has some risks:

- Placing heat-generating light sources inside an incubator disrupts the temperature and humidity regulation inside the incubator by providing a localized point source of heat not under control of the incubator environmental feedback controller.
- Electronics placed inside the warm, humid and acidic environment of a cell culture incubator can suffer accelerated wear.
- Microscopes made from components that are not suitable for extended exposure to incubator conditions can experience mechanical, structural, or electronic failures.

While existing incubator microscopes purport to address these challenges, methods used to enclose and protect light sources and electronics inside the incubator also generally make these instruments inflexible. We will present a design which circumvents this problem entirely by minimizing the amount of sensitive components inside the incubator.

Cell plate “hotel” systems designed for high content microscopy address both of these issues by having the microscope outside of an incubator, but also automating the process of transferring and then returning the cell plate to an incubator with a robotic motion system^17,18,24^. This minimizes the amount of time that a dish spends outside of an incubator and allows for high-throughput automation of experiments. However, these systems are restricted to use with specific form factors of cell culture plates to maintain compatibility with the external incubation system and the automated motion system. Furthermore, the change in environment and mechanical agitation of transferring plates to and from the microscope causes physiological perturbations which are avoided in incubator-integrated imaging solutions. Due to the need for an external incubation system and robotic manipulators these systems physically occupy a large amount of space^25^. The high cost of these systems and makes these devices difficult to deploy in academic laboratory settings.

Each of these system classes exhibits intrinsic limitations that constrain the adoption of live-cell microscopy by availability, experimental throughput, temporal resolution and imaging duration. Biological specimens undergo highly dynamic, time-dependent morphological and phenotypic changes, such that high-frequency, long-term live-cell microscopy will capture transient behaviors, cell–cell interactions, and temporal patterns of signaling or differentiation otherwise missed in single imaging timepoints. The limited accessibility of automated, high-resolution longitudinal imaging platforms therefore imposes a bottleneck on experiments involving biological tissues.

To help mitigate the costs of commercial microscopy systems and improve accessibility of specialized microscopy experiments, individual laboratories are developing their own specialized scientific equipment^26^. This approach has been an important part of the growing field of open laboratory hardware: scientific laboratory equipment designed to be assembled by end-user laboratories and whose designs, components, and fabrication methods are made publicly available for publication under an open license^27^. Many pieces of laboratory equipment have open hardware designs, including centrifuges^28^, pipettes^29^, pumps^30^, and more complex instruments like sampling systems for bioreactors^31^.

Advancements in open hardware techniques have allowed for the rapid development and deployment of many bespoke microscopy systems. These range from handheld resolution enhancers like Cybulski et al.’s Foldscope^32^, to full-size experimental systems like Ly et al.’s Picroscope^33^ and Collins et al.’s OpenFlexure modular microscope framework^34^. There are also open hardware microscopes for fluorescence microscopy, like Wincott et al.’s Microscopi, Zehrer et al.’s UC-STORM and Stewart and Gianni’s OPN Scope^35–37^. However, the vast majority of these microscopes are not designed for live cell imaging in the hot and humid environment of a cell culture incubator, which is usually kept at 37°C and near-saturated humidity for mammalian cell culture. This elevated temperature causes permanent warping of 3D printed components due to heat creep, exposes sensitive electronics to moisture and corrosion, and presents thermal management challenges with electronics and light sources^33^. Furthermore, the humid environment promotes microbial growth, porous 3D printed materials resist surface decontamination, and most 3D printed materials, especially those printable in a standard consumer setting, are not autoclave-stable. To this end, the device we report will also serve as a demonstration of the benefits of using machined metals in open hardware for incubator use. We take advantage of CNC milling and routing prototyping service providers to manufacture robust metal optomechanical and structural components suitable for long-term use in an incubator. This allows open hardware to be constructed from components which are stable to incubator temperatures, nonporous and easily heat-sterilizable for antisepsis, and durable and corrosion-resistant. Our mechanically robust, purpose-engineered design produces a microscopy platform capable of continuous, long-term operation under standard incubator conditions.

A common shortcoming of incubator microscopes is that components such as light sources or controllers can be damaged by the incubator or cause damage to the samples in the incubator^2^. Moving these components outside the incubator allows the microscope to continuously capture images for weeks at a time inside an incubator without perturbing the biology in the incubator [Figure 3a].

In an effort to make live time-series microscopy, we present a modular, compact, and automated multi-fluorescence microscope that supports imaging with up to four channels and that can be operated at incubator temperatures for extended periods with a high imaging frequency of up to one image per second, exposure time permitting. Our device automates image acquisition and has a modular and easily reconfigurable design, allowing form factor, light sources and inclusion of plate scanning automation to be changed as needed on an experiment-by-experiment basis. We demonstrate versions of this microscope with multi-fluorescence, with plate-scanning automation and image stitching, and with integration with fluidic automation.

## RESULTS

### 1.1 System Design

This microscope [Figure 1] is designed for continuously performing high frequency imaging of living samples over weeks inside a cell culture incubator, while maintaining device functionality and minimizing impact on the incubator environment. Light sources, power supplies, and controllers are placed outside the incubator to reduce thermal load, preserve internal volume, and protect both electronics and the sample from excess heat and humidity. Illumination is delivered via a fiber optic cable routed through the rear incubator port. An illustration of how the microscope is kept separate from the light sources is shown in Figure 1b, where a cutaway shows how a fiber optic cable is run through a port on the back of the incubator. The modularity of this system enables live imaging of both conventional cell culture plates and complex experimental systems such as bioreactors, microfluidic circuits, and other larger devices. Additionally, using fiber optic cables as inputs allows for any type of light source that has an output terminal that can be converted to a SMA905 thread to be used as an input.

**Figure 1:**
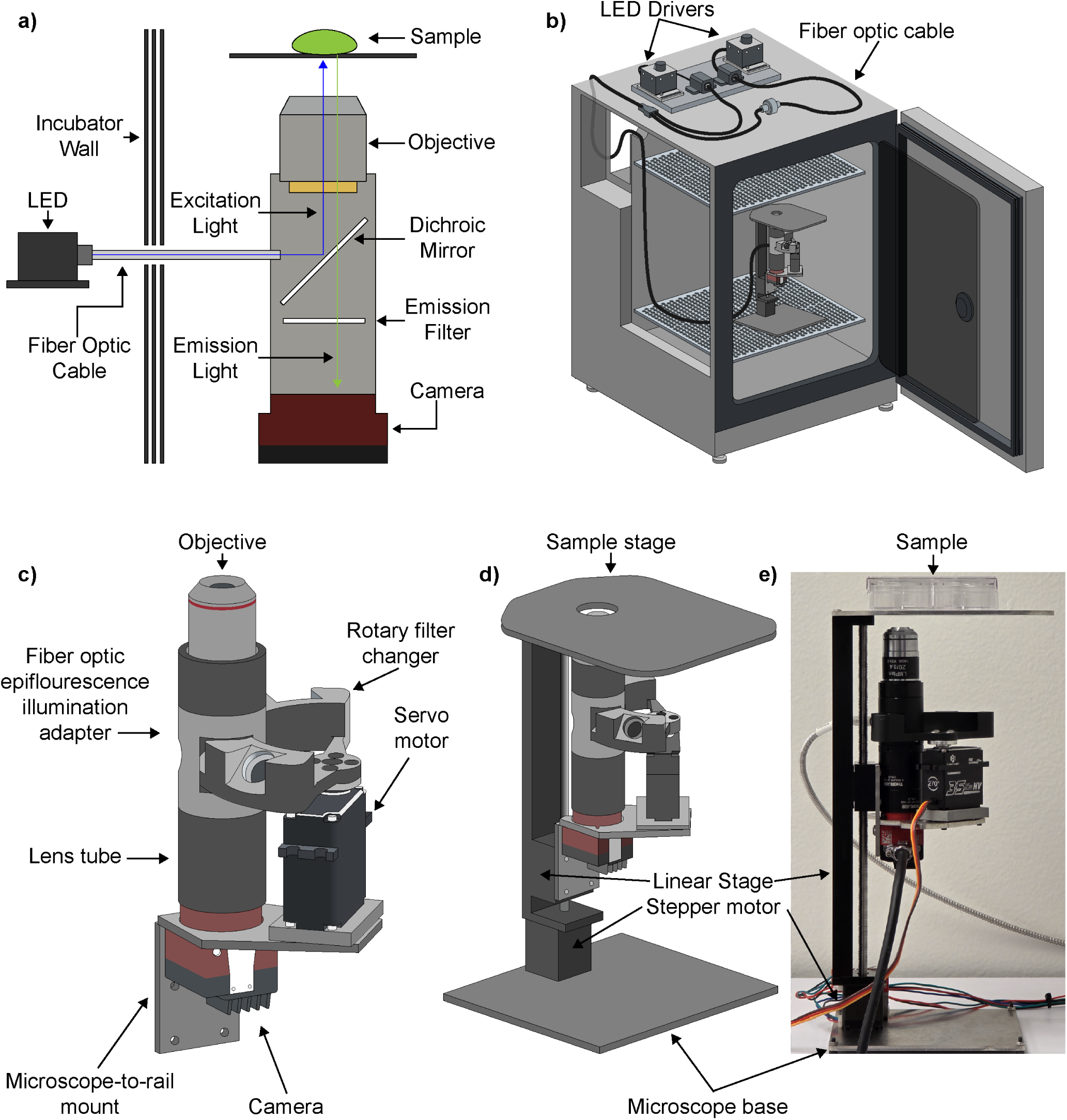
Design of modular in-incubator multi-fluorescence microscope. a) Diagram of excitation and emission light paths through the microscope. b) Render of the microscope inside the cell culture incubator connected to its external illumination system. c) Render of optical components of the microscope. d) Render of the assembled microscope, including vertical motion system and frame. e) Photograph of the microscope deployed on a laboratory bench.

For fluorescence imaging, dichroic mirrors and emission filters are mounted on a motorized rotary filter changer turret [Figure 1c], which aligns the desired filter set with the light being emitted from the end of the fiber optic cable [Figure 1a]. This setup enables multi-fluorescence imaging in our fiber optic remote illumination design.

The device also has the ability to perform routines like z-stacking or the creation of extended depth of field composites. This is accomplished through the use of a computer controlled vertical axis motion system for remotely or automatically controlling how close the objective is to the sample. The use of a Z-axis rail for mounting also allows for the use of objectives with varied working distances. A render and photo of the system mounted to the Z axis motion system can be seen in Figure 1d and Figure 1e.

To ensure structural durability under elevated temperature and humidity, consumer-grade 3D printing materials, particularly plastics, commonly used in open hardware, were excluded due to heat creep, which causes deformation and misalignment over time. This allows the device to operate for extended periods (days to weeks) either inside and outside a cell culture incubator. The microscope is built from stainless steel and aluminum, metals which resist corrosion in the warm, humid, acidic incubator environment. These materials were also selected due to their compatibility with the device sterilization process described in Section 1.6; individual metal parts can also be autoclaved. Metal parts for this system comprise sheet metal fabrications, off-the-shelf commercial metal optical components and metal CNC milled components.

A device built to be used in cell culture incubators should be designed to take up as little space as possible, since incubator space is a limited resource. Thus, the form factor of the microscope has been kept to approximately 150mm x 120mm x 230mm, which allows four microscopes to be on one shelf inside a standard 150 to 170-liter cell culture incubator. Other in-incubator microscopes have a larger footprint that still can fit cleanly inside of a typical 150 to 170 liter incubator. In comparison, Merces et al.’s Incubot occupies a 300mm × 370mm × 370mm volume, and Etaluma’s LS820 has dimensions of 240mm x 226mm D x 278mm^15,38^.

#### 1.1.1 Optical Design

While the Allied Vision Alvium 1800 U-319m USB and 1800 U-291m cameras used in this device are advertised as being intended for use in machine vision applications, the pixel size, quantum efficiency and finely controllable exposure make these units suitable for biological fluorescence imaging. The overall cost and small physical footprint make the camera module used here an affordable and more compact alternative to sCMOS cameras used in high end microscopy equipment. Its compact size also makes it a sensible choice for use inside of an incubator, since most conventional sCMOS cameras are significantly larger and have additional cooling requirements. Because of their shared form factor and software interface, any camera from the Allied Vision Alvium 1800 line with a C-Mount thread can be used with this microscope, but the 319m and 291m were the only ones tested in the experiments presented in this manuscript, and were chosen for their particular specifications, as described in Section 1.4.

Synchronizing the excitatory light and the capture of images to limit unnecessary exposure, which could cause phototoxicity, was accomplished using software and hardware components described in 1.4.

#### 1.1.2 Assembly Subunits

The microscope consists of three main modules: the frame, optical assembly, and external lighting system [Figure 1a]. These modules can be easily exchanged with other subsystems of the same type used in the microscope variants presented in this paper. All of our frame, optics, and lighting modules are designed to be cross-compatible to allow for mix-and-match flexibility in creating microscopy systems for specific experimental designs.

The frame includes the base, linear rail and sample stage. Both the base and the sample stage are laser cut from stainless steel. The imaging surface has a circular opening large enough to accommodate the objective’s field of view but small enough to support 35 mm dishes without risk of falling through. Any plate, slide, or sample with a transparent bottom and within the working distance of the objective can be imaged this way.

The optical assembly consists of the camera [Section 1.4], lens tube [Figure 1c], objective [Figure 1c], rotary filter system [Figure 1c], dichroic mirror[Figure 1a], and emission filter [Figure 1a] (specifications in Section 1.4). The camera is connected to the microscope objective by a series of C-mount lens tubes and optical adapters [Figure 1c]. A custom black anodized aluminum epifluorescence illumination adapter within this optical path integrates the fiber optic input and supports the rotary filter changer [Figure 1c]. This adapter has a tight-fitting curved slot that allows smooth filter rotation while blocking stray excitatory light.

The rotary filter changer holds up to four filter sets, each consisting of a 12.5 mm dichroic mirror and emission filter, secured with threaded rings [Figure 4b]. It is driven by a coreless servo motor mounted to a stainless steel bracket that connects back to the camera module [Figure 1c]. A 7mm thick piece of milled aluminum acts as a spacer between the bracket and the motor to position it in the plane of the filter rotation module. Above the adapter, a C-mount lens tube holds the microscope objective, connected via an RMS-to-C-mount adapter. The entire assembly is mounted to a vertical linear actuator using a stainless steel bracket [Figure 1c]. A diagram of the optical path, showing excitatory and emission light, is provided in [Figure 1a].

The external lighting system is modular and scalable, consisting at a minimum of an LED, an LED driver, and a fiber optic cable delivering light into the incubator [Figure 1b]. Light intensity is controlled by a microcontroller connected to a computer. For multi-color experiments, bifurcated fiber optic bundles can combine multiple LEDs into a single output beam. Each LED has its own driver, with all sources controlled by a single microcontroller [Figure 1b]. Component specifications and control protocols are detailed in Section 1.4. The system is compatible with standard optical breadboards for mounting, and components can be positioned anywhere within the fiber cable’s reach, typically on or beside the incubator [Figure 1b]. This system could also be made to be compatible with 2mm bore or greater light guides that have SMA905 connectors or smooth ferrule-in-collar attachments for which threaded adapters exist to integrate into our system.

#### 1.1.3 Two Axis Scanning Stage Module

To enhance the capabilities of this microscope while still maintaining a flexible form factor, we designed a module to allow for X and Y axis motion that consists of two additional linear actuators installed in a larger frame that can be installed by replacing the base of the microscope. A rendering of this system can be seen in Figure 6a. This system allows for scanning regions of interest in a sample anywhere inside of a 86mm x 128 mm space, which corresponds with common dimensions of standard 6, 12, and 24 well cell culture plates.

### 1.2 Microscope Design Validation

#### 1.2.1 Thermal Validation

To measure the effects of the microscope components on the ambient temperature of the incubator, and the temperature stability of important microscope electrical components, adhesive thermocouples were placed inside of a cell culture dish on the imaging surface of the microscope, in the incubator air space beside the microscope, on the body of the camera module, and on the servo motor operating the rotary filter changer. The readings from these thermocouples were taken for more than 40 hours while camera and motor were moved every two minutes to match the conditions of the experiment presented in Section 1.3.2.

These readings showed that the localized temperature around each thermocouple was stable to within +/-0.1 degrees Celsius after an initial equilibration period, and that even though there were areas with higher local temperatures (like the body of the camera), the temperature inside the empty cell culture dish and the ambient temperature of the incubator were not perturbed from their physiological setpoint. The inside of the well plate read at 36.8°C, the empty space next to the microscope read at 37.4°C, the body of the camera read at 46.4°C, and the body of the servo motor read at 41.2°C. A chart showing a representation of this data can be seen in Figures 3c. These values are well below the maximum operating temperatures of the camera and servo, which are 65°C and 60°C, respectively.

Like other live cell microscopes, this device needs to adjust to the thermal environment of the incubator for a short time to prevent undesired focal plane changes due to thermal expansion^39^. We found that two hours was enough time to allow the microscope to adjust to the incubator conditions, with no focal plane drift observed after this point. This is required only for initial placement into a heated cell culture incubator; subsequent experiments do not need this prerequisite as long as the incubator remains heated and the device remains inside.

#### 1.2.2 Resolution Validation

The high fidelity of our optical system design is sufficient for all features on a USAF 1951 resolution target slide (Edmund Optics) to be distinguishable at 10x or 20x magnification (up to group 9-3), giving this microscope the ability to resolve features as small 1.55***µ***m (645 line pairs per millimeter).

#### 1.2.3 Positioning Repeatability

The linear actuator for controlling vertical positioning is driven by a 1.8 degree per step stepper motor using a leadscrew with a 1mm pitch. Without microstepping, this gives a theoretical 5 micron resolution in Z-axis positioning. An endstop was mounted to the bottom of the actuator to provide a consistent reference zero point for positioning, and to mitigate the effects of backlash. For the two axis scanning stage module, the two additional axes consist of a pair of linear rails with travel lengths of 150mm and 100mm respectively. These rails are driven in the same manner as that of the microscope itself, and are rated to support up to 15kg in vertical load. These rails are driven by a 1.8 degree per step stepper motor using a leadscrew with a 5mm pitch. Without microstepping, this gives a theoretical 25 micron resolution in X and Y-axis positioning. An endstop was mounted to the side of both actuators to provide a consistent reference zero point for positioning, and to mitigate the effects of backlash. Any imprecision in the motion of these axes can be mitigated through the use of capturing overlapping images and then stitching them together.

### 1.3 Biological Imaging Experiments

To demonstrate the in incubator longitudinal imaging capabilities of the microscope, we conducted representative experiments that highlight the two most important features of this microscope; that it can capture images for extended periods inside a cell culture incubator, and that it can perform multi-fluorescence imaging tasks while doing so.

#### 1.3.1 Long Term *in vitro* longitudinal live imaging of a human iPSC derived vascular organoid

To highlight the stability of our system in its intended use environment, a 30 day old human iPSC derived vascular organoid expressing TdTomato was imaged every 3 minutes for approximately 14 days without interruption, providing 6714 images in total. This organoid was maintained using a variant of microfluidic system based on the device described by Seiler et al. and Voitiuk et al.^40,41^. As shown in Figure 3 and discussed further below, continuous changes in cell morphology were observed in the tissue over the duration of the experiment. A subset of the resulting images from this experiment with labeled time points are shown in Figure 3. A timelapse movie generated from the images captured during the experiment can be seen in the repository linked in the data availability statement.

##### Analysis

Our apparatus allowed us to observe growth and remodeling dynamics of the vascular organoid. Vascular organoids undergo sprouting, rearrangement of cells, recruitment of various cell types including endothelial cells and pericytes to self-assemble and form capillary network^42^. At day 0 and day 1.0, the organoid shows smooth edges around its border; as early as day 1.5 of embedding in the GelMA matrix, radial sprouting of cells is observed around the edges and further uniform extension of this pattern progresses until the end point of the experiment. This demonstrates the vessel sprouting when embedded in the GelMA matrix. Vascular network remodeling is another feature in self-organizing vascular organoids^43^. Formation of microcapillary-like structures is visible as early as day 0 (timepoint D0 in Figure 3a) and they continue to grow, branch, and thicken on day 13 (timepoint D13.6 in Figure 3a). In addition, the re-arrangements of these microcapillaries can be observed from day 5 images as far as these images show that we observe the self-organizing vascular organoid, thickening of the microcapillary vascular cells and branching points and the increase in overall size.

#### 1.3.2 Multi-fluorescence *in vitro* Imaging of co-culture of primary human brain microvascular endothelial cells and pericytes

To demonstrate the multi-fluorescence capabilities and high temporal imaging frequency of this device inside the incubator, a well containing a 2:1 ratio of endothelial cells tagged with eGFP and pericytes tagged with the fluorophore mCherry was observed for approximately 3 days.

Images were captured on both fluorescent channels in pairs using a 20x long working distance LM Plan objective (Boli Optics), with 2 minutes between each pair of image captures. Dual color imaging was accomplished through the use of the rotary filter changer described in Section 1.4 and shown in Figures 1 and 4a and the multiple input light source scheme described in Section 1.4 and seen in 4b. This experiment captured 2266 two color timepoints over the course of three days without interruptions. Images were postprocessed and composited using the method described in Section 1.14. A representative subset of images captured during this experiment can be seen in 4c.

##### Analysis

Endothelial cells form the innermost lining of the blood vessel and serve functions in vascular health, nutrient and oxygen delivery, and homeostasis^44^. Pericytes serve as supporting mural cells covering the endothelial cells to provide tight junction points. Together these cells have a key role in maintaining blood-brain barrier^45^. We seeded endothelial cells (ECs) and pericytes at 2:1 ratio at sub confluent levels and captured time-lapse imaging of these cells. The time series shows cell division, migration and recruitment of pericytes by ECs. ECs form invasive tip cells that are similar to filopodia and are migratory in nature [Figure 3 H15.9]. From time lapse we could observe endothelial cells recruiting pericytes to their vicinity and once recruited, pericytes wrap around the abluminal surface of endothelial cells. One pericyte covers several endothelial cells. Comparison of the endothelial cell area and number of connected components suggests that our microscopy experiment demonstrates endothelial networks coalescing.

#### 1.3.3 Microscope Scanning of Human Brain Organoid

To demonstrate the modularity of the system and a whole and the efficacy of the two axis attachment described above in Section 1.1.3, still images and directional scans were taken of a KOLF 2.2J human induced pluripotent stem cell (hiPSC) derived dorsal forebrain organoid in which some of the neurons are expressing GFP via Adeno-Associated Virus introduction of a hSyn-GFP reporter cassette that drives GFP expression specifically in neurons^46^. Due to the low emission of the sample, a LumeDEL N490 LED with a maximal output of 140mW was paired with a 1500um bore multimode fiber optic patch cable with a numeric aperture of 0.5 (Thorlabs M107L01). The output from the patch cable was then passed through a collimator (LumeDEL CL12G) to maximize the amount of light being transported to the sample. The collimator was mounted to the side of a 30mm filter cube (Thorlabs CM1-DCH) containing an excitation filter (Chroma Technologies ET480/30x) and a rectangular dichroic mirror (Edmund Optics 69-899). The design of this system and results captured with it can be seen in Figures 6a, 6b, and 6c. X-Y axis scans were captured by using the coordinates of the respective linear rail to calculate a 30% horizontal overlap between adjacent images, with a vertical stack being taken at each of these positions. These images were then processed according to the method shown in Figure 6d and Section 1.14.

##### Analysis

HiPSC dorsal forebrain organoids recapitulate key aspects of human specific cortical development that cannot be otherwise modeled. At 22 weeks, the organoids contain a mixture of excitatory and inhibitory neurons, neural and glial progenitors, and astrocytes. We observe circular patches of green, GFP-labeled neurons at the outer surface of the organoid [Figure 6b-c]. The zoomed-in image at the bottom of Figure 6b demonstrates that neurons (labeled in green) within these cortical organoids are capable of generating intricate networks of dendrites and axons replicating the structure and function of mature neurons in the developing cortex. By this time point at week 22, most of the excitatory neurons of the cortical plate have already formed, and these neurons begin to generate connections via neurites portrayed by the arrows on the bottom of Figure 6b.

## DISCUSSION

We have demonstrated a modular platform for capturing long term longitudinal image datasets of live fluorescent biological samples inside of a cell culture incubator. By developing a modular incubator-compatible microscope, we hope to lower the barrier to entry for groups to perform long-term time series imaging experiments inside incubators or incubator-like environments by giving them the flexibility to customize the microscope to meet their own needs. Our tests with a variety of biological samples and device configurations demonstrate that our system can function effectively inside of an incubator for extended periods to generate high-frequency fluorescence imaging datasets.

This was accomplished through our design and manufacturing approach which improves on the current open hardware norm of plastic-based desktop 3D printing, instead using rapid prototyped metal components. We believe that our device remains accessible and open; while many labs lack in-house metalworking capabilities, rapid prototyping service bureaus offer small-run custom part fabrication at a cost competitive with in-house plastic 3D printing when accounting for scientist labor, equipment and supplies. Alternatively, these parts can be manufactured inhouse at institutions with multi-axis CNC milling capabilities. We hope that our approach might serve as a template for others to combine robust manufacturing with open labware approaches to create bespoke, built for purpose tools especially those designed for humid and thermally intensive or otherwise challenging environments.

Looking forward, our system has potential for use in many types of experiments not covered in our demonstrations. The modular nature of our microscope design allows the straightforward addition of other imaging modalities to demonstrate more complex applications of this device. These could include capturing the differentiation of cells that have cell type specific reporters, performing lineage tracing, tracking cells with exceptionally high motility, and using z-stacks to gather three dimensional multi-fluorescent imaging datasets at each time point^6,47^. Other applications of interest could be tracking the transient expression of proteins^48^, live cell biochemical dye assays such as viability dyes, hypoxia indicators or metabolism specific probes, and functional fluorescence imaging tasks like calcium imaging^49^. We also envision that more modules can easily be integrated into our system, like a objective changer, a phase contrast system, or support for the use of lasers and other non-LED light sources. These additions would provide more utility and flexibility in the design of custom microscopy systems using our modular framework.

## METHODS

### 1.4 Optics

To detect of fluorophores emitting in the the green range like eGFP, a 500nm dichroic mirror (Edmund Optics 69-866) was paired with a 525nm/50nm Full Width at Half Maximum (FWHM) emission filter (Chroma Technologies ET525/50m). The excitatory light used was either a 455nm peak emission fiber-coupled LED (Thorlabs M455F3) or one of two 490nm peak emission fiber-coupled LEDs (LumeDEL N490 or Thorlabs M490F4). For the 490nm fiber-coupled LEDs, a 480nm/30nm FWHM excitation filter (Chroma Technologies ET480/30x) was used. Out of the these three sources, the LumeDEL N490 fiber-coupled LED has the highest output power (140mW compared to 17mW for the 455nm LED and 1.8mW for the other 490nm LED, respectively).

To detect fluorophores emitting in the red range like tdTomato, a 600nm dichroic mirror (Edmund Optics 69-868) was paired with a 630nm/75nm FWHM emission filter (Chroma Technologies ET630/75m). The excitatory light used was a 554nm peak emission fiber-coupled LED (Thorlabs MINTF4). A 560nm/40nm FWHM excitation filter (Chroma Technologies ET560/40x) was used.

To detect fluorophores emitting in the blue range like DAPI, a 400nm dichroic mirror (Edmund Optics, 69-864) was paired with a 460nm/50nm FWHM emission filter (Chroma Technologies AT460/50nm). The excitatory light used was a 375nm fiber-coupled LED (Thorlabs M375F3). A 375nm/28nm FWHM excitation filter (Chroma Technologies ET375/28x) was used.

While any fluorophore, LED, and filter combination will work with this device, fluorophores in the green, red and blue ranges were tested and validated in this study. We tested the device with both live and fixed samples.

This microscope is equipped with a custom filter switching module, which allows it to image multiple fluorophores. This filter switching module consists of a custom C-mount threaded barrel with a curved slot cut into it that allows for a rotating turret block that holds dichroic mirrors and emission filters to slide through to change which filter set is in the optical path of the microscope and excitation light port. This filter holding turret accommodates up to 4 pairs of dichroic mirrors and emission filters. A coreless servo motor (Flash Hobby M35CHW) is used to control the angular position of the filter holder relative to the C-mount barrel, positioning filters and dichroic mirrors in the optical path. The choice of a servo motor allows for positioning precision and repeatability. This approach compresses the bulky form factor of filter cubes and allows for the use of multiple filter sets in a smaller space. While the assembly is a multi-filter monobloc design, the filters are of standard circular 12.5 mm format installed with threaded retaining rings and can be easily changed.

For single channel fluorescence imaging, in order to deliver excitation light from the light remote source to the apparatus, a step-index multimode fiber optic patch cable with a 1500 micron bore and a numerical aperture of 0.5 (Thorlabs M107L01) was used to direct excitatory light into the filter assembly. To deliver multiple sources of excitatory light into one optical path, SMA905 multimode fiber bundles (Thorlabs BFY1000LS02) are used to direct multiple possible input light sources. The system is compatible with any LED fiber optic coupled light sources with SMA905 connectors, and these light sources can be paired with multi-mode fiber optic cables and bundles of any length and bore size. An illustration of the light path showing both excitatory and emissive light is shown in Figure 1a.

For multi-fluorescence imaging, a 1000***µ***m bore bifurcated SMA905 fiber bundle (Thorlabs BFY1000LS02) is mounted so the split ends are each connected to an LED while the shared end connects to the barrel of the microscope, allowing multiple light sources to be connected to one output point. This arrangement is shown in Figure 1b. Fiber coupling used SMA905 format adapters (Thorlabs).

The shared end of the input fiber bundle (regardless of the number of inputs) is connected to the custom filter switching barrel to be lined up with the dichroic mirror to allow for the use of multiple bands of excitatory light in one experiment. This excitatory light can also be passed through a collimator (LumeDEL CL12G or Thorlabs F950SMA-A) to reduce its spread within the optical pathway, to improve effective sample illumination power.

Each light source is a fiber-mounted LED. LEDs from Thorlabs are driven by a T-Cube LED driver (Thorlabs LEDD1B), and LumdeDEL LEDs use an integrated driver. To modulate the intensities of light sources that were controlled by the T-Cube LED driver, an analog signal was provided by an FTDI FT232H USB breakout board (Adafruit 2264) connected to a digital to analog converter (DAC) with an MCP4725 chip (Sparkfun BOB12918) using the I^2^C communication protocol. Enabling and disabling LEDs is done automatically with the DACs connected to each light source being modulated by the FT232H breakout board. For light sources that use an integrated driver like the LumeDEL N490, the intensity of the light was modulated using the Python pyserial library with over a RS232 cable that ships with the light source using settings provided by the manufacturer in their documentation.

### 1.5 Data Acquisition and Microscope Control

Images were captured with one of two camera modules:

- Allied Vision Alvium 1800 U-319m USB camera, which uses a Sony IMX265 3.2MP CMOS monochrome global shutter sensor with a maximum image resolution of 2064 x 1544 pixels, and a pixel size of 3.45***µ***m x 3.45***µ***m.
- Allied Vision Alvium 1800 U-291m USB Camera, which uses a Sony IMX421 3.2MP CMOS monochrome global shutter sensor with a maximum image resolution of 1944 x 1472 pixels, and a pixel size of 4.5***µ***m x 4.5***µ***m.

Any camera from the Alvium 1800U series of USB cameras from Allied Vision will be directly plug-and-play compatible with this system, but these two specific modules were chosen due to having a balance of high quantum efficiency, large pixel size, and relatively low cost (approximately $700).

We chose a camera module with a C-mount thread, which allows the use of commercial lens tubes and optical hardware. C-mount extension barrels and adapters (Thorlabs) were used with custom components to mount fiber optics, dichroic mirrors, and emission filters [Figure 1]. A C-Mount to RMS adapter (Thorlabs RMSA5) allows for the use of standard microscope objectives with this system. In the images presented here, 4x (Edmund Optics 59-934), 10x (Edmund Optics 59-935), and 20x (Boli Optics MT05073431) objectives were used, and their respective fields of view when used with our system are shown in Figure 2a.

**Figure 2:**
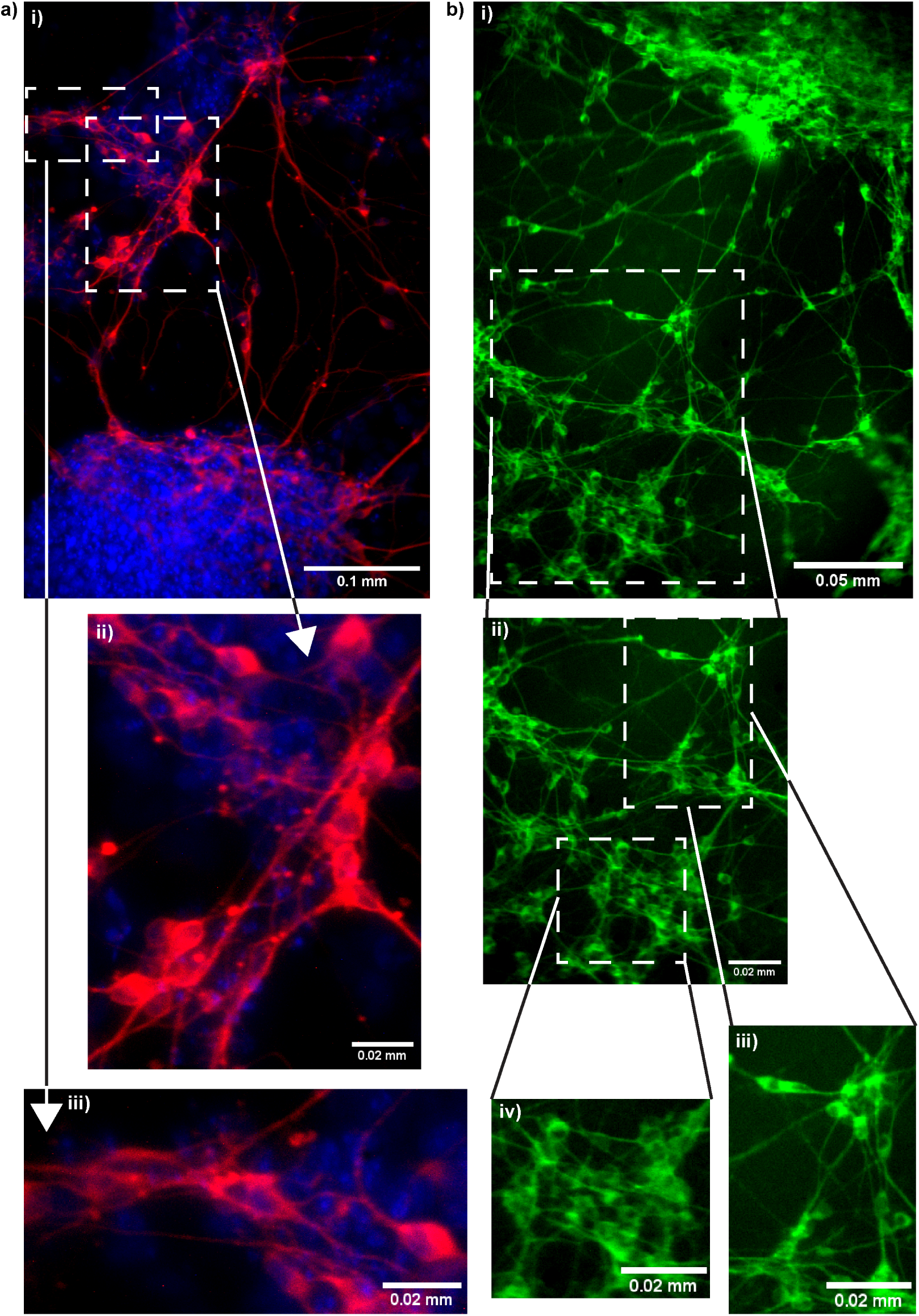
Immunohistochemistry (IHC) images. a) i) IHC image of mouse neurons. ii) Neurons expressing MAP2+ are tagged with Alexa Fluor 546 (red), and nuclei are stained with DAPI (blue). Neuronal somas with radiating MAP2+ dendrites. iii) Soma dendritic junctions. b) i) IHC image of mouse neurons expressing MAP2 tagged with Alexa Fluor 488 (green). The image shows a more spread-out neuoronal network. ii), iii), and iv) densely clustered group of neurons with closely apposed somas and extensive dendritic arborization. MAP2+ processes extend radially from the soma, forming a complex and highly interconnected dendritic network. The neurons exhibits multiple dendritic projections, suggesting active neurite outgrowth and synaptic connectivity. This aggregate morphology is characteristic of maturing neuronal cultures and reflects a network formation in vitro.

**Figure 3:**
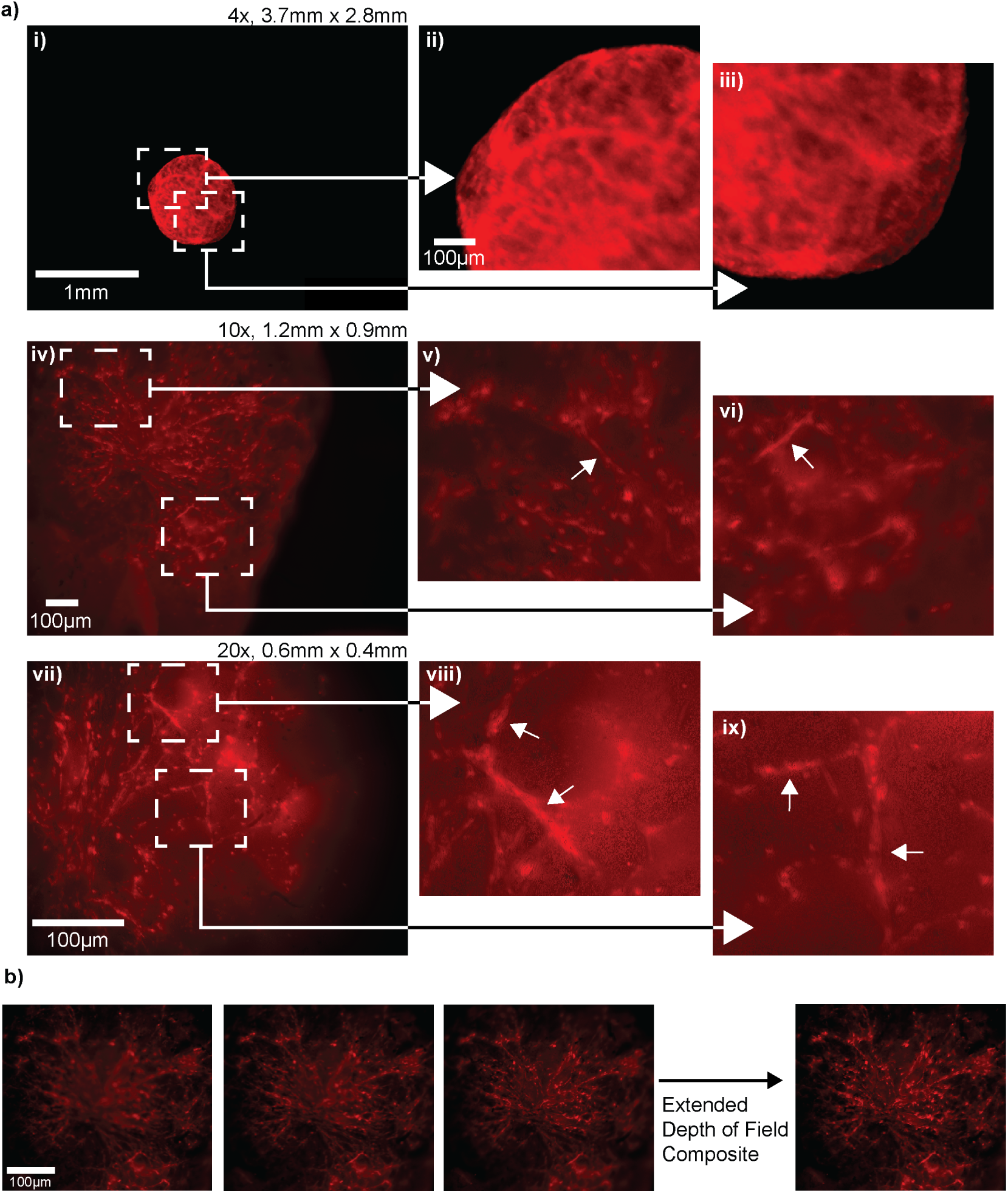
Image Resolution and Compositing. a) tdTomato (red) tagged vascular organoids at varying magnifications, denoting the real world size of the fields of view (4x objective −3.7mm x 2.8 mm). 10x and 20x are extended depth of field composites. In the inset images, the 3D organoid surface exhibits overlapping network from the 3D vascular organoid into overlapping network that coalesce into denser vascular structure emerging from the surface (top). The surface of the vascular organoid is forming vascular network that are radially branching on the surface of the organoid. The arrows point to a capillary like elongated and curved structure that is distinct from the surrounding cells. A vascular capillary structure is connecting to other microvessel like structure on both ends (middle). The vascular organoid has an extended capillary network with branching points are clearly visible in the image. The vascular organoid surface depicts capillaries with bifurcation points and several branching points along the path of the developing capillary network, these vessels are larger than the surrounding capillaries as they develop mimicking human brain capillary network (bottom).

Images were acquired from the camera using the Python implementation of OpenCV and the Python library vmbpy (provided by Allied Vision for use with their camera modules), which allows control over exposure time and gain. A rail-based linear actuator with a T6x1mm lead screw (ZeberoXYZ) was used for Z-axis positioning control and was driven by a Pololu Tic N<u>°</u> T825 stepper motor controller using its manufacturer provided command line API, ticcmd. This actuator was fitted with a mechanical switch endstop to provide more consistent positioning, using the inbuilt endstop functionality of the motor controller. Using our provided example programs in the repository for this document (https://github.com/braingeneers/incubatormicroscopesupplement) as a template, users can create their own custom imaging routines to best suit their needs.

### 1.6 Device and Sample Preparation

To decontaminate the microscope for use in an aseptic incubator environment, all surfaces of the device and cables connected to it were wiped down with sporocidal Oxivir Tb Disinfectant Cleaner wipes (Diversey 4599516) and sprayed with 70% ethanol or isopropanol solution. The device was then placed inside the incubator, and cables for the camera, linear rail, servo motor and fiber optic were run through the port in the back of the incubator and sealed with a pass-through incubator plug to minimize leakage of CO_2_.

### 1.7 Primary cell culture

Human brain microvascular endothelial cells (ECs) (ScienCell 1000) and human brain vascular pericytes (PCs) (ScienCell 1200) were cultured in endothelial culture medium (ScienCell 1001) supplemented with 5% fetal bovine serum (Sciencell 0025), 1% Endothelial Cell Growth Supplement (Sciencell 1052) and 1% penicillin/streptomycin (Sciencell 0503) and pericyte medium (ScienCell 1200) supplemented with 2% fetal bovine serum, 1% Pericyte Growth Supplement (Sciencell 1252) and 1% penicillin/streptomycin solution as per manufacturer’s instructions. Cells were passaged at 70% confluency. Cells were maintained in a humidified 5% CO_2_ incubator.

### 1.8 Labeling of primary cells

ECs and PCs were labeled with green fluorescent protein (GFP) and mCherry, respectively. Passage 2 cells underwent lentiviral transduction with either EF1a-GFP or CMV-mCherry (1 *×* 10^8^) (VectorBuilder). Cells were treated at an MOI of 5 for 24 h. Expression of the fluorescent markers was confirmed by fluorescence microscopy after 2-3 days. The cells were subselected for GFP and mCherry expression using flow cytometry and expanded for further use in experiments.

### 1.9 Two color imaging experiment of primary cells

Briefly 5 *×* 10^5^ GFP-endothelial cells and 2.5 *×* 10^5^ mCherry-pericytes (EC:PC-2:1) were seeded in a 24-well cell culture plate, allowed to attach for 2 days. Cells were at passage number 5. Cells were washed with PBS and replenished with fresh culture media at the start of the experiment. Images were acquired for 3 days every 2 minutes using a 554nm fiber-coupled LED (Thorlabs MINTF4) to excite the red channel and a 455nm fiber-coupled LED (Thorlabs M455F3) at full intensity to excite the green channel, using the filtersets described in Section 1.4.

### 1.10 Human induced pluripotent stem cell culture

Human induced pluripotent WTC11tdTomato stem cells were a gift from the Nowakowski lab (University of California, San Francisco). hiPSCs were cultured in a Matrigel (Corning 354277) coated plate with mTESR1 Plus media (Stemcell Technologies 100-0276) changed every day. Cells were passaged with ReLeSR (Stemcell Technologies 100-0484). Accutase (Sigma-Aldrich A6964) was used to generate a cell suspension for organoid aggregation.

### 1.11 Vascular organoid generation

Vascular organoids were generated from hiPSCs following Wimmer et al.’s protocol^42^. Briefly, hiPSCs were detached with Accutase and 12,000 cells were seeded per well of an ultra-low attachment 96-well plate with 10 µM of ROCK inhibitor Y-27632 (10 µM) (Tocris Bioscience 1254). After 24 hours, a complete media change was performed into mesoderm induction media consisting of N2B27 medium supplemented with CHIR99021 (12 µM) (Tocris Bioscience 4423) and BMP-4 (2 µM) (Miltenyi Biotech 130-111-164) for 3 days. Subsequently, media was changed to vascular induction media consisting of N2B27 media supplemented with VEGF-A (100ng/ml) (Peprotech 100-20) and forskolin (2 µM) (Sigma-Aldrich F3917), and maintained for another 3 days until vascular sprouts were observed^42^. These resulting vascular organoids were individually embedded in 30 µl droplets of Matrigel on silicone organoid aggregation sheets (Stemcell 08579) on ice and then incubated at 37°C for 1 hour to allow solidification of the Matrigel around the vascular organoid. Embedded organoids were transferred to ultra-low attachment 6-well plates and cultured in vascular organoid maintenance media consisting of Stempro-34 media (Gibco 10640) supplemented with VEGF-A (100 ng/ml), FGF-2 (100 ng/ml) (Stemcell Technologies 78003) and 15% fetal bovine serum (Gibco A525680) and maintained on an orbital shaker at 90 rpm until further use, with media changes every 2 days.

### 1.12 Imaging experiment for vascular organoid

A day 30 matrigel-encapsulated vascular organoid was embedded in 5% gelatin-methacrylate (GelMA) (Cellink IKG125000005) which was crosslinked with 0.5% Lithium phenyl-2,4,6-trimethylb (Cellink 5269). In a 24 well plate, 200 µl of GelMA was poured and overlaid with a 30 day old vascular organoid and overlaid on top with 200 µl of GelMA. The GelMA was then UV crosslinked for 30 sec and 500 µl of Stempro-34 media supplemented with VEGF-A (100 ng/ml) (Peprotech 100-20) and FGF-2 (100 ng/ml) (Stemcell Technologies 78003). Media was changed every 6 h using an automated microfluidics unit^40,41^. Every 2-3 days, the media in the reservoir connected to the pre-programmed automatic microfluidics pump [Section 1.13] was exchanged with VEGF-A and FGF-2 supplemented fresh media.The vascular organoid was imaged for 2 weeks, over which time images were captured every 3 min. Excitatory light was supplied by a 554nm fiber-coupled LED (Thorlabs MINTF4) using the filterset described in Section 1.4.

### 1.13 Automated media exchange during imaging

Cell culture media exchange was performed by an automated culturing system^40,41^ scheduled to exchange 250 ***µ***L of vascular organoid maintenance media every 6 hours. The fresh media was drawn from a refrigerated reservoir, allowed to warm to room temperature in line and delivered via FEP tubing (Cole Parmer AAD02119-CP) to a custom 3D-printed fluidic insert. The tubing was mated to the insert using PEEK chromatography compression fittings sealed with PTFE tape. The 3D-printed fluidic insert, adapted to fit a standard 24-well plate, featured inlet and outlet ports positioned for optimal media addition and removal retention as described previously^40,41^. No lid was used; instead, a silicone O-ring (McMaster Carr 5233T297) held the insert in place. This and a Breathe-Eazy microcentrifuge tube sealing film sealed the well while allowing gas exchange.

A fluidic insert and volume-occupying ring to center the organoid were printed in Formlabs Biomed Clear V1 biocompatible resin using a Form 3B+ SLA printer. The print was post-processed with Form Wash and Form Cure devices according to the manufacturer’s instructions. 3D printed parts were sterilized by autoclave, then assembled. After assembly, the wetted parts of the system were sterilized using Cidex OPA ortho-phthaldehyde solution, rinsed with PBS and primed with media.

### 1.14 Microscopy Image Correction Methods

Across all experiments, post-experimental image adjustments were performed done using FIJI (FIJI is Just ImageJ) software50.

Single color images were loaded either alone or as part of a sequence and then had a lookup table (LUT) applied to them using the inbuilt ImageJ “LUT” tool (preset “Green”, “Red”, or “Blue” depending on the fluorophore used), and then had their contrast adjusted using the “Brightness/Contrast” tool. For adjusting sets or sequences of images, they were converted to an image stack to allow for uniform batch adjustment. When needed, speckles in the image created due to a high gain level were removed using the “Despeckle” tool.

Multi-color still images were created by using the above protocol for separate images acquired per channel, followed by compositing using the “Merge Channels” tool. The two channels of the image were then aligned manually using the “Align RGB Planes” plugin.

Vertical stacks were created using the “Extended Depth of Field” plugin in easy mode51.

Image sequences were registered to each other in bulk using the “Linear Stack Alignment with SIFT” tool, and image stitching was performed using the “Pairwise Stitching” tool52,53.

To perform image deconvolution to further improve contrast, a point spread function (PSF) was generated using the “PSF Generator” plugin using the Born and Wolf model^54,55^. PSFs were generated based on the peak wavelength, immersion medium, numerical aperture, and the pixel size of the imaging system used. The input image and the PSF were then passed into the “DeconvolutionLab2” plugin using the Richardson-Lucy algorithm for 10 iterations^56–58^.

### 1.15 Image Registration and Alignment

To correct for mechanical drift and ensure spatial consistency in longitudinal datasets, we implemented a custom image registration pipeline using the Enhanced Correlation Coefficient (ECC) algorithm^59^. The registration logic was developed in Python utilizing the OpenCV library^60^.

To enable the visual comparison of experimental runs and the manual validation of channel registration, we developed an interactive Video Overlay and Aligner tool using the OpenCV framework^60^. This software overlays two synchronized or independent video streams (for example, two different fluorescent channels or time-lapse sessions) and applies real-time geometric transformations to achieve spatial alignment. For longitudinal datasets where samples may shift or grow non-uniformly over time, the software supports a keyframe-based alignment workflow.

### 1.16 User Centered Design Methods

When creating custom laboratory hardware like this microscope, it is important to consider the needs of the user^61^. In the creation of scientific instruments like our microscopy system, the adoption of rapid prototyping methodologies emerges as a relevant and practical user-centered design approach. Rapid prototyping involves quickly iterating on previous designs, integrating feedback on minor revisions to build towards a finished product, and is particularily popular in the open hardware space^62,63^. In this application, expert level scientists from multiple scientific disciplines have been frequently consulted and collaborated with the development and testing of this device as a part of our rapid prototyping cycles. These collaborations included for testing the microscope on a wide range of sample types, determining relevant hardware and software features, and assisting in the design of modules for bespoke experiments which can be seen in other works featuring other versions of our system^64–67^.

## RESOURCE AVAILABILITY

### Lead contact

Requests for further information and resources should be directed to and will be fulfilled by the lead contact, Yohei Rosen.

### Materials availability

This study did not generate new materials.

### Data and code availability

Bills of material, assembly guides, module comparison charts, and wiring diagrams can be found in the supplemental material.

2D and 3D part files, microscope control software, thermal validation data, and a timelapse movie for the data shown in Figure 4 can be found in a Git repository located at https://github.com/braingeneers/incubatormicroscopesupplement.

**Figure 4:**
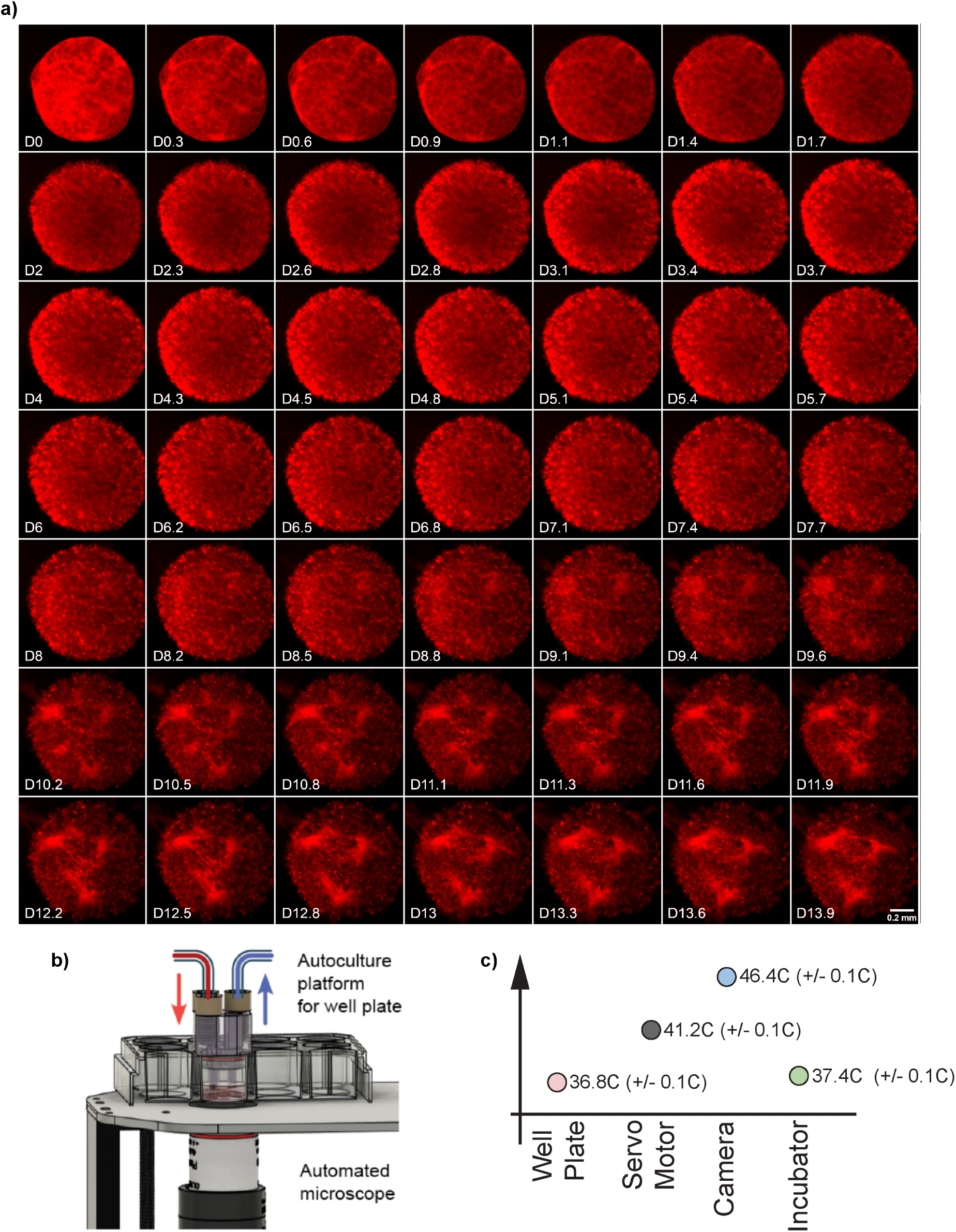
Longitudinal study of tdTomato vascular organoid. a) Grid of images showing the growth of the organoid over the period of 14 days. b) Illustration of the automated fluidics system used in Section 1.12. c) Temperature chart for the quantification experiment described in Section 1.2.1, where temperatures were monitored for 40 hours. Stated range represents the entire range of captured values after thermocouple readings stabilized 2 hours into the experiment.

**Figure 5:**
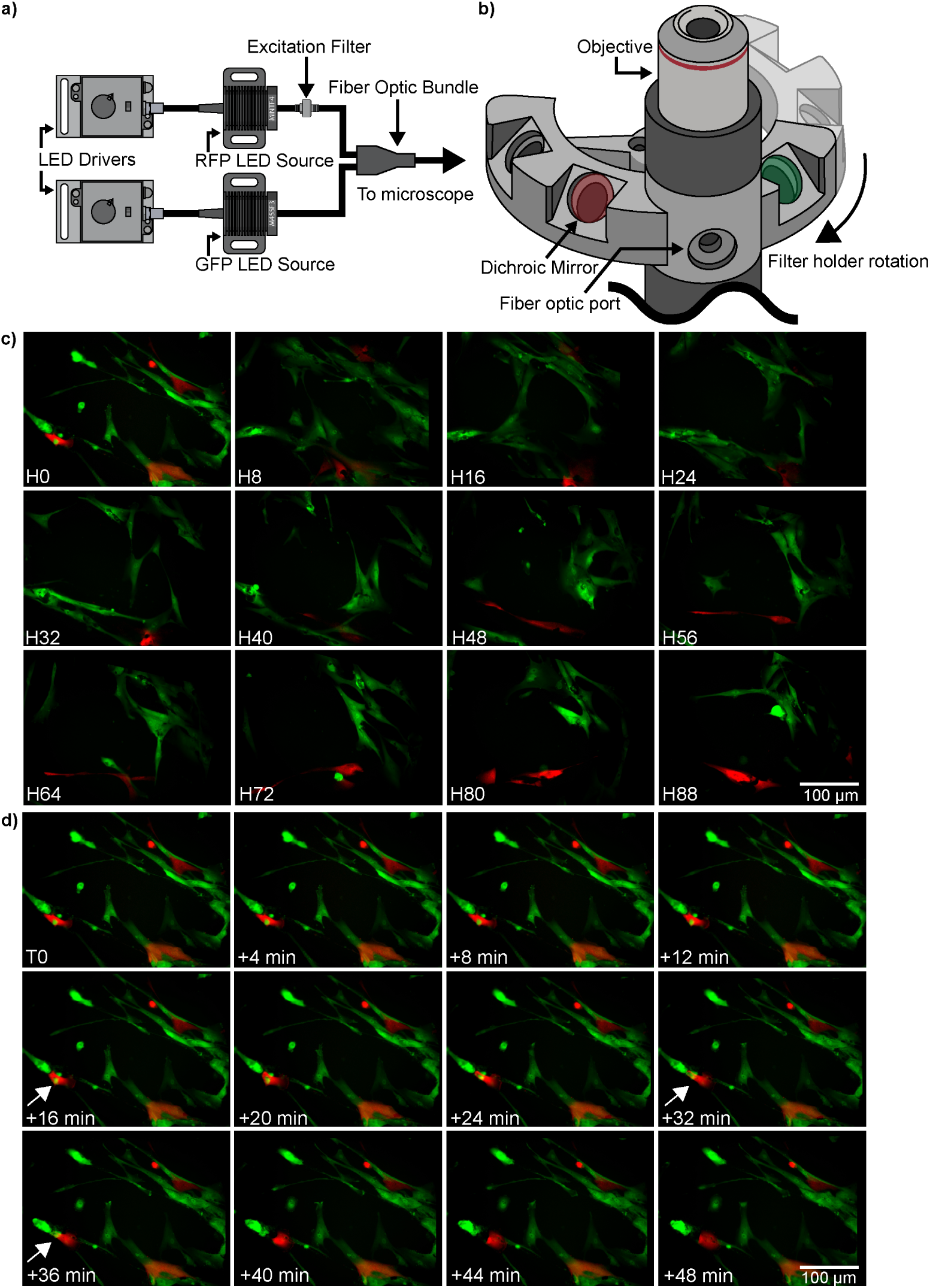
Multi-fluorescence observation of different vascular cell types. a) Diagram showing arrangement of optical components for two-color multi-fluorescence imaging with endothelial cells (green) and pericytes (red). b) Illustration of rotary filter changer. c) Grid of images subsampled to multi-hour intervals to demonstrate superficial changes in morphology of the sample over the course of 4 days. d) Grid of images showing a subset of images from the experiment (starting from H0) at 4 minute intervals. The microscope demonstrates the temporal and spatial resolution required to study cell divisions and cell-cell interactions. As example, from T0-T+36 min we observe two dividing endothelial cells (green dots: nuclei of GFP-ECs, white arrow head). The divided cells subsequently express cytoplasmic GFP and align to the pericyte surface (T+32) are stabilized by the pericyte at (T+36). Two channel images were aligned using the method described in 1.15

**Figure 6:**
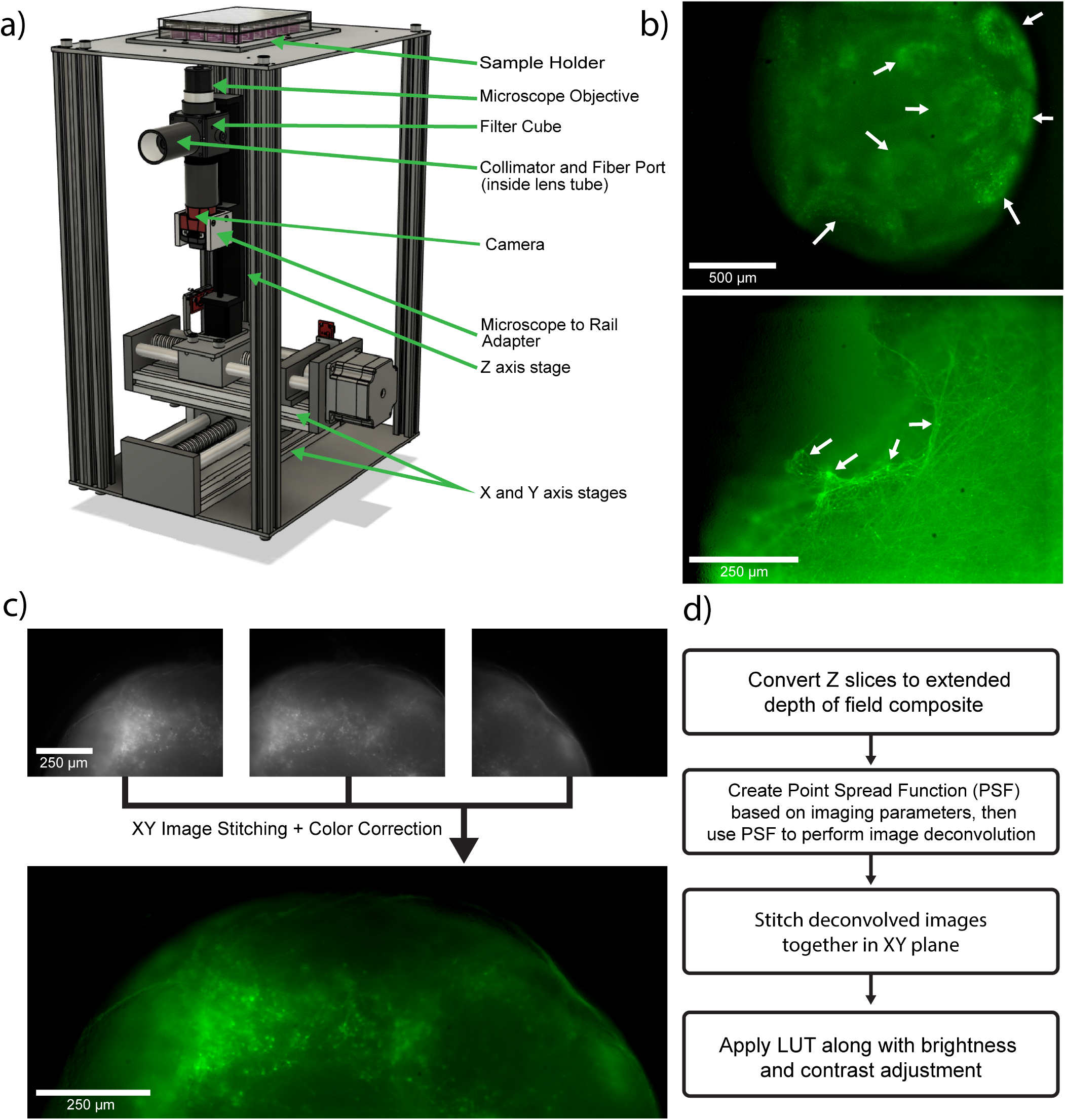
Design and implementation of three axis motion system. a) Diagram showing components of one-color fluorescence three axis microscope. b) Images of 22 week old dorsal forebrain organoids captured using LumeDEL N490 LED source. In the top image, the arrows point to neural rosette-like structures. In the bottom image, the arrows point to neurites connecting adjacent neurons. c) A composite image of a dorsal forebrain organoid consisting of 3 source vertical stacks in adjacent horizontal positions. d) Flow chart that describes the process of combining horizontally adjacent vertical stacks into an image like that seen in c).

- Bills of material, assembly guides, module comparison charts, and wiring diagrams can be found in the supplemental material. All other data reported in this paper will be shared by the lead contact upon request.
- 2D and 3D part files, microscope control software, thermal validation data, and a time-lapse movie for the data shown in Figure 4 can be found in a Git repository located at https://github.com/braingeneers/incubatormicroscopesupplement.
- Any additional information required to reanalyze the data reported in this paper is available from the lead contact upon request.

## Supporting information

Supplemental Documentation

## ACKNOWLEDGMENTS

This work was supported by the Schmidt Futures Foundation under award SF857 (S.R.S., D.H. and M.T.), the National Human Genome Research Institute under award RM1HG011543 (S.R.S, D.H., S.K. and M.T.), the National Science Foundation under awards 2134955 (S.R.S, D.H. and M.T.), 2034037 (M.T.) and 2515389 (M.A.M.-R., D.H. and M.T.), the National Institute of Mental Health under award U24MH132628 (M.A.M.-R. and D.H.), the National Institute of Neurological Disorders and Stroke under award U24NS146314 (M.A.M.-R. and D.H.), the California Institute for Regenerative Medicine under awards DISC4-16285 (M.A.M.-R., S.R.S. and M.T.) and DISC4-16337 (M.A.M.-R.), the University of California Office of the President under award M25PR9045 (M.A.M.-R., S.R.S. and M.T.), the Brain & Behavior Research Foundation under award 33184 (M.A.M.-R.), and the University of California Santa Cruz Center for Information Technology Research in the Interest of Society and the Banatao Institute Interdisciplinary and Innovation Program (I2P) (M.A.M.-R). Y.R. was supported by a CIRM postdoctoral fellowship under Award number CIRM EDUC4-12759. J.L.S and M.M. were supported by the UCSC IRACDA fellowship program under National Institutes of Health (NIH) award number K12GM139185. S.V.-C. was partially supported by the Graduate Pedagogy Fellowship from the University of California Santa Cruz Teaching and Learning Center.

We also thank Dr. Pierre Baudin, Dr. Victoria Ly, Valeska Victoria, Hunter Schweiger, Max McGlynn, Dr. Nico Hawthorne, Ryan Fenimore, Dr. Matthew Elliott, and the rest of the Braingeneers both past and present at the UCSC Genomics Institute for their invaluable support on this project over the past four years. We thank Jeff Jackson for support during the patent disclosure process, and the support engineers at Thorlabs, Edmund Optics, Chroma Technologies, and Allied Vision Technologies for their encyclopedic knowledge and advice regarding their catalogs.

## AUTHOR CONTRIBUTIONS

D.E. and Y.R. designed and built the device. Y.R. conceived of the device and the study. D.E. drafted the manuscript. D.E and Y.R. revised the manuscript. S.A. provided and maintained the blood vessel organoids and microvascular cells shown in the results section, and wrote sections related to the experiments conducted with them. S.V-C, K.D., M.M, and J.S. provided biological samples for the testing and validation of the device. K.V. and S.S. provided an automated fluidics system for the blood vessel organoid experiment. J.M. designed and built the computer vision pipeline and performed data analysis on the two color primary cells and vascular organoid datasets. A.T. authored figures, graphics, and renders. S.N. provided foundational support in creating initial device designs. S.K., M.A.M.-R, E.W., D.H., S.R.S, and M.T. provided supervision and laboratory resources. During the preparation of this work, the authors used ChatGPT and Claude to improve clarity and sentence structure. Github Copilot was also used for the development of imaging analysis pipelines. After using this tool/service, the authors reviewed and edited the content as needed and assume full responsibility for the content of the publication.

## DECLARATION OF GENERATIVE AI AND AI-ASSISTED TECHNOLOGIES

During the preparation of this work the authors used ChatGPT to edit text for conciseness. LLM-based coding assistants were used to assist with python scripting specifically in data processing and visualization workflows. After using these tools, the authors reviewed and edited the content as needed and take full responsibility for the content of the published article.

## SUPPLEMENTAL INFORMATION INDEX

Notes S1-S4 and their legends in a PDF

