## Supplemental Documentation for "A Modular In-Incubator Microscope for Longitudinal Live Cell Microscopy"

### 1 Supplementary Note 1: Device Comparisons

#### Fiber Optic Cable Comparison Chart

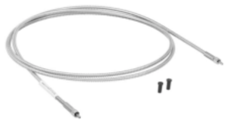

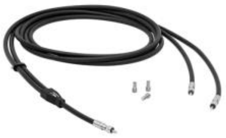

| | Multimode Fiber Optic Patch Cable | 2-to-1 Multimode Fiber Optic Patch Cable Bundle |
|---|---|---|
| Bore Size | Ø1500um | Ø1000um (Ø500um per end) |
| Numeric Aperture | 0.5 | 0.39 |
| Catalog Number | Thorlabs M107L02 | Thorlabs BFY1000LS02 |

#### CMOS Camera Comparison Chart

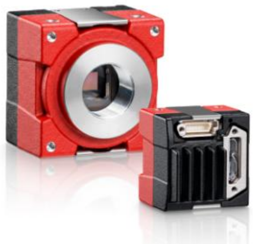

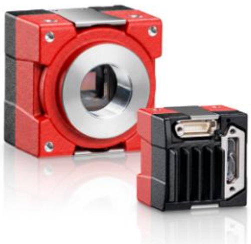

| | Allied Vision Alvium 1800U-291m USB Camera | Allied Vision Alvium 1800U-319m USB Camera |
|---|---|---|
| Camera Sensor | Sony IMX421 | Sony IMX265 |
| Pixel Size | 4.5µm x 4.5µm | 3.45µm x 3.45µm |
| Image Dimensions | 1,944 × 1,472 pixels | 2,064 x 1,544 pixels |
| Sensor Format | 2/3 | 1/1.8” |

#### Light Source Comparison Chart

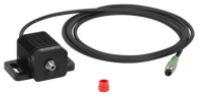

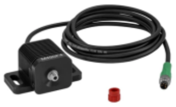

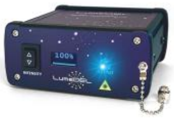

| | Thorlabs M455F3 | Thorlabs M490F4 | LumeDEL N490 |
|---|---|---|---|
| Output Power (typical rating, from manufacturer) | 24.5mW | 2.8mW | 100mW |
| Peak Wavelength | 455nm | 490nm | 495nm |
| Bandwidth (Full Width Half Max) | 14nm | 26nm | 20nm |

### Optics Core Comparison Chart

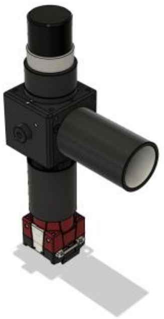

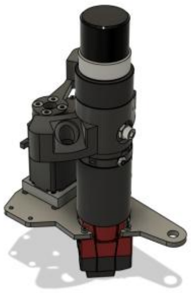

| | Filter Cube | Filter Turret |
|---|---|---|
| Fluorophores Supported | 1 | Up to 4 |
| Collimator Support | Yes | No |

### Microscope Frame Comparison Chart

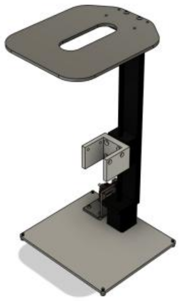

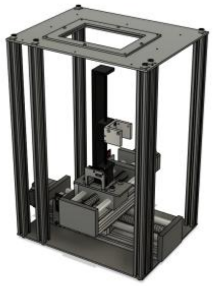

| | One Axis Frame | XYZ Motion System |
|---|---|---|
| Axes of Motion | 1 | 3 |
| Travel | 150mm in Z | 150mm in X, 100mm in Y, 100mm in Z |
| Repeatability Precision | +/- 5µm | +/- 3µm in X and Y, 5µm in Z |

### 2 Supplementary Note 2: Ordering Guides

#### 2.1 Z Frame Bill of Material

| *Identifier* | *Part Name/Number OR File Location* | *Source* | *Qty.* | *Cost Per* | *Material* |
|---|---|---|---|---|---|
| **Electronics** | | | | | |
| Linear Rail | Zeberoxyz 150mm Travel Mini Linear Rail | Amazon | 1 | 69.89 | |
| Linear Rail Driver | Tic T825 Stepper Motor Driver | Polulu | 1 | 42.95 | |
| Endstop Switch | Mechanical Endstop Limit Switch Module (Red) | Amazon | 1 | 8.99 | |
| 330Ω Resistor | 330 ohm Resistor 1/2W | Amazon | 1 | 4.59 | |
| Linear Rail Power Supply | ALITOVE 24V 2A Power Supply | Amazon | 1 | 15 | |
| **Custom Parts** | | | | | |
| Microscope Base | /CAD/flatparts/microscope_base_z.dxf | SendCutSend | 1 | 20 | Stainless Steel (304 series) (.125") |
| Microscope Top | /CAD/flatparts/flatslot_top_oval.dxf | SendCutSend | 1 | 20 | Stainless Steel (304 series) (.125") |
| Camera Holder | /CAD/CNC/U_rail to alvium adapter.stp | PCBWay | 1 | 30 | Stainless Steel 304 (CNC) |
| Endstop Holder | /CAD/flatparts/endstopholder_Z.dxf | SendCutSend | 1 | 7 | Stainless Steel (304 series) (.125") with 90 DEGREE BEND |
| **Fasteners** | | | | | |
| M2.5x25 Socket Head Screws | 91290A055 | McMaster Carr | 1 | 9.5 | |
| M3 Socket Head Screw Set | M3x6-M3x30 Socket Head Screws Kit | Amazon | 1 | 7 | |
| M3x8 Countersinking Screws | 92125A128 | McMaster Carr | 1 | 6.42 | |
| M3 Button Head Screw Set | M3x6-M3x30 Button Head Screws Kit | Amazon | 1 | 8 | |
| *Cost basis for all listed parts in in USD as of January 2026.* | | | | | |
| *Files for all items in the custom parts section can be found at the github repository linked in the main manuscript.* | | | | | |

#### 2.2 XYZ Frame Bill of Material

| *Identifier* | *Part Name/Number OR File Location* | *Source* | *Qty.* | *Cost Per* | *Material* |
|---|---|---|---|---|---|
| **Electronics** | | | | | |
| Z Axis Linear Rail | Zeberoxyz 1PCS Mini Linear Rail, 150mm Effective Stroke Length | Amazon | 1 | 65.89 | |
| Bottom Axis Linear Rail | Zeberoxyz 1PCS 150mm Length Dual Optical Axis Guide Ballscrew SFU1605 | Amazon | 1 | 100.8 | |
| Middle Axis Linear Rail | Zeberoxyz 1PCS 100mm Length Dual Optical Axis Guide Ballscrew SFU1605 | Amazon | 1 | 95.8 | |
| Endstop Switch | Mechanical Endstop Limit Switch Module (Red) | Amazon | 3 | 8.99 | |
| 330Ω Resistor | 330 ohm Resistor 1/2W | Amazon | 3 | 4.59 | |
| Power Supply for all Three Axes | 24V 10A Power Supply Adapter, 100-240V AC to 24V 10A | Amazon | 1 | 30.59 | |
| Linear Rail Driver | Tic T825 Stepper Motor Driver | Polulu | 3 | 42.95 | |
| **Custom Parts** | | | | | |
| Camera Holder | /CAD/CNC/U_rail to alvium adapter.stp | PCBWay | 1 | 30 | Stainless Steel 304 (CNC) |
| Endstop Hitter for Bottom Axis | /CAD/flatparts/endstop_hitter_long_xyz.dxf | SendCutSend | 1 | 6 | 6061 T6 Aluminum (.375") |
| Endstop Hitter for Middle Axis | /CAD/flatparts/endstophitter_short_xyz.dxf | SendCutSend | 1 | 12 | 6061 T6 Aluminum (.375") |
| Endstop Holder | /CAD/flatparts/endstopholder_XYZ.dxf | SendCutSend | 2 | 8 | 6061 T6 Aluminum (.125") |
| Bottom to Middle Axis Adapter | /CAD/flatparts/midbot_countersink_xyz.dxf | SendCutSend | 1 | 23 | 6061 T6 Aluminum (.125") |
| Middle to Top Axis Adapter | /CAD/flatparts/midtop_longer_countersink_xyz.dxf | SendCutSend | 1 | 16 | 6061 T6 Aluminum (.125") |
| Z Endstop Holder | /CAD/flatparts/xyz_end_218_rev.dxf | SendCutSend | 1 | 7 | Stainless Steel (304 series) (.125") |
| Microscope Top | /CAD/flatparts/xyz_frame_top_turner.dwg | SendCutSend | 1 | 70 | Stainless Steel (304 series) (.125") |
| Microscope Bottom | /CAD/flatparts/bigbase_xyz.dxf | SendCutSend | 1 | 77 | Stainless Steel (304 series) (.125") |
| Plate Holder | /CAD/flatparts/frame_top_xyz.dwg | SendCutSend | 1 | 10 | 5052 H32 Aluminum (.063") |
| **Mounting** | | | | | |
| 400mm 2020 Aluminum Extrusions | 4pcs 400mm T Slot 2020 Aluminum Extrusion | Amazon | 2 | 19 | |
| 2020 Aluminum Extrusion Slot Covers | 6 Meter 1010 2020 Aluminum Extrusion T Slot Cover | Amazon | 1 | 12 | |
| **Fasteners** | | Amazon | 1 | 10 | |
| M6 Socket Head Screw Kit | M6 Screws with Nuts Assortment Kit Hex Socket Head Cap Bolts | Amazon | 1 | 10 | |
| M5 Socket Head Screw Kit | M4 Screws with Nuts Assortment Kit Hex Socket Head Cap Bolts | Amazon | 1 | 10 | |
| M4 Socket Head Screw Kit | M5 Screws with Nuts Assortment Kit Hex Socket Head Cap Bolts | Amazon | 1 | 10 | |
| M2.5 Socket Head Screw Kit | M2.5 Screws with Nuts Assortment Kit Hex Socket Head Cap Bolts | Amazon | 1 | 10 | |
| M3 Flathead Countersinking Screw Kit | M3 Countersunk Screws Bolts Nuts Washers Assortment Kit | Amazon | 1 | 10 | |
| M3 Button Cap Screw Kit | M3 Screw Assortment Kit, M3 Screws Hex Button Head Cap Screws Bolts Nuts | Amazon | 1 | 10 | |
| M2.5x32 Flathead Screws | iexcell 100 Pcs M2.5 x 32mm Thread Pitch 0.45 mm Stainless Steel 304 Hex Socket | Amazon | 1 | 10 | |
| *Cost basis for all listed parts in in USD as of January 2026.* | | | | | |
| *Files for all items in the custom parts section can be found at the github repository linked in the main manuscript.* | | | | | |

### 2.3 1 Color Optics Core Bill of Material

| *Identifier* | *Part Name/Number OR File Location* | *Source* | *Qty.* | *Cost Per* |
|---|---|---|---|---|
| **Optics** | | | | |
| Filter Cube | CM1-DCH | Thorlabs | 1 | 205.91 |
| Dichroic Mirror | 500nm, 25.2 x 35.6mm, Dichroic Longpass Filter (cat 69-899) | Edmund Optics | 1 | 192 |
| Double Male C-Mount Adapter | C-Mount Double Male Rotating Ring (cat 53-865) | Edmund Optics | 1 | 77.5 |
| SM05 to C-Mount Adapter | SM05A2 | Thorlabs | 1 | 24.65 |
| SM05 Lens Barrel w/ Retaining Ring | SM05L03 | Thorlabs | 1 | 15.96 |
| C-Mount Lens Barrel | CML40 | Thorlabs | 1 | 31.79 |
| Collimator | Collimator Lens Diameter 12.5 mm, Molded glass NA 0.63 w/ SMA Connector (cat 70-920) | Edmund Optics | 1 | 260 |
| SM1 to C-Mount Adapter | SM1A9 | Thorlabs | 1 | 22.88 |
| C-Mount to SM1 Adapter | SM1A10 | Thorlabs | 1 | 23.82 |
| 4x Microscope Objective | 4X Nikon Achromatic Finite Conjugate Objective (cat 59-934) | Edmund Optics | 1 | 83 |
| 10x Microscope Objective | 10X Nikon Achromatic Finite Conjugate Objective (cat 59-935) | Edmund Optics | 1 | 123 |
| 20x Microscope Objective | 20x Long Working Distance LM Plan Achromatic Metallurgical Microscope Objective Lens | Boli Optics | 1 | 228.98 |
| SM1 Lens tube w/ Retaining Ring | SM1L20 | Thorlabs | 1 | 19.1 |
| GFP Excitation Filter | ET480/30x (25mm Circle Size) | Chroma Technologies | 1 | 365 |
| GFP Emission Filter | ET525/50m (12.5mm Circle Size) | Chroma Technologies | 1 | 365 |
| SM1 Cap | SM1CP2 | Thorlabs | 1 | 22.13 |
| 15mm C-Mount Lens Tube | CML15 | Thorlabs | 1 | 22.93 |
| C-Mount to RMS Adapter | RMSA5 | Thorlabs | 1 | 23.68 |
| USB Camera | Allied Vision Alvium 1800U-319m Right Angle (cat 23-204) | Edmund Optics | 1 | 604.95 |
| USB Camera Cable | AVT Type-A to Micro-B, USB 3.0 Locking Cable, 3m (cat 14-903) | Edmund Optics | 1 | 123 |
| *Cost basis for all listed parts in in USD as of January 2026.* | | | | |
| *Files for all items in the custom parts section can be found at the github repository linked in the main manuscript.* | | | | |

### 2.4 2 Color Optics Core Bill of Material

| *Identifier* | *Part Name/Number OR File Location* | *Source* | *Qty.* | *Cost Per* | *Material* | |
|---|---|---|---|---|---|---|
| **Optics** | | | | | | |
| USB Camera | Allied Vision Alvium 1800U-319m Right Angle (cat 23-204) | Edmund Optics | 1 | 604.95 | | |
| USB Camera Cable | AVT Type-A to Micro-B, USB 3.0 Locking Cable, 3m (cat 14-903) | Edmund Optics | 1 | 123 | | |
| Dichroic mirror for RFP | 600nm, 12.5mm Diameter, Dichroic Longpass Filter (cat. 69-868) | Edmund Optics | 1 | 143 | | |
| RFP Emission Filter | ET630/75m (12.5mm Circle Size) | Chroma Technologies | 1 | 365 | | |
| Dichroic mirror for GFP | 500nm, 12.5mm Diameter, Dichroic Longpass Filter (cat 69-866) | Edmund Optics | 1 | 143 | | |
| GFP Emission Filter | ET525/50m (12.5mm Circle Size) | Chroma Technologies | 1 | 365 | | |
| Dichroic mirror for DAPI | 400nm, 12.5mm Diameter, Dichroic Longpass Filter (cat 69-864) | Edmund Optics | 1 | 143 | | |
| DAPI Emission Filter | AT460/50m (12.5mm Circle Size) | Chroma Technologies | 1 | 225 | | |
| 40mm C-Mount Lens Tube | CML40 | Thorlabs | 1 | 31.79 | | |
| Fiber Mounting Point | HASMA | Thorlabs | 1 | 10.04 | | |
| Filter Retaining Ring | SM05RR | Thorlabs | 1 per filter | 4.63 ea | | |
| 15mm C-Mount Lens Tube | CML15 | Thorlabs | 1 | 22.93 | | |
| C-Mount to RMS Adapter | RMSA5 | Thorlabs | 1 | 23.68 | | |
| 4x Microscope Objective | 4X Nikon Achromatic Finite Conjugate Objective (cat 59-934) | Edmund Optics | 1 | 83 | | |
| 10x Microscope Objective | 10X Nikon Achromatic Finite Conjugate Objective (cat 59-935) | Edmund Optics | 1 | 123 | | |
| 20x Microscope Objective | MT05073431 20x Long Working Distance Objective | Boli Optics | 1 | 228.98 | | |
| **Electronics** | | | | | | |
| Servo Motor | M35CHW Coreless Servo | Flash Hobby | 1 | 41.99 | | |
| Servo Driver | Micro Maestro 6-Channel USB Servo Controller | Pololu | 1 | 27.95 | | |
| **Custom Parts** | | | | | | |
| Camera Servo Mount | /CAD/flatparts/cam_to_turner_servo.dwg | SendCutSend | 1 | 12 | Stainless Steel (304 series) (.125") | |
| Servo Booster Seat | /CAD/CNC/booster7mm.step | PCBWay | 1 | 30 | SLM Aluminum (3D Printed) | |
| C-Mount to Turner Adapter | /CAD/CNC/rotator to c-mount.stp | PCBWay | 1 | 37 | Aluminum 6061 | Black Anodizing + Bead Blast |
| Turner Turret Bottom | /CAD/CNC/rotary_filter_barrel_bottom.stp | PCBWay | 1 | 33 | Aluminum 6061 | Black Anodizing + Bead Blast |
| Turner Turret Top | /CAD/CNC/turret_outer_top.stp | PCBWay | 1 | 37 | Aluminum 6061 | Black Anodizing + Bead Blast |
| Turner Turret | /CAD/CNC/rotaryinner.step | PCBWay | 1 | 37 | Aluminum 6061 | Black Anodizing + Bead Blast |
| **Fasteners** | | | | | | |
| M2 Screw Kit | M2 Socket Head Screws/Nuts Kit M2x4mm-M2x20mm | Amazon | 1 | 8 | | |
| M3 Screw Kit | M3 Button Head Screws/Nuts Kit M3x6mm-M3x30mm | Amazon | 1 | 8 | | |
| M3 Countersinking Screw Kit | M3 Flat Head Screws/Nuts Kit M3x4mm-M3x20mm | Amazon | 1 | 6 | | |
| *Cost basis for all listed parts in in USD as of January 2026.* | | | | | | |
| *Files for all items in the custom parts section can be found at the github repository linked in the main manuscript.* | | | | | | |

#### 2.5 Thorlabs LED and Fiber Optics Bill of Material

| *Identifier* | *Part Name/Number OR File Locat* | *Source* | *Qty.* | *Cost Per* |
|---|---|---|---|---|
| **One Color Green** | | | | |
| 490nm LED | M490F4 | Thorlabs | Choose this or 455nm | 377 |
| 455nm LED | M455F3 | Thorlabs | Choose this or 490nm | 502.1 |
| Fiber Optic Patch Cable | M107L02 | Thorlabs | 2 | 260.18 |
| GFP Excitation Filter | ET480/30x (12mm Circle Size) | Chroma Technologies | 1 | 365 |
| Excitation Filter Holder Part 1 | SM05SMA | Thorlabs | 1 | 35.59 |
| Excitation Filter Holder Part 2 | PM20-SMA | Thorlabs | 1 | 40.36 |
| LED Driver | LEDD1B | Thorlabs | 1 | 380.04 |
| LED Driver Power Cable | KPS201 | Thorlabs | 1 | 43.15 |
| Microcontroller for LED Control | FT232H | Adafruit | 1 | |
| Digital to Analog Converter | BOB-12918 | Sparkfun | 1 | |
| BNC Male Plug Terminal Block | 2888 | Adafruit | 1 | 0.95 |
| **One Color Red** | | | | |
| Mint LED | MINTF4 | Thorlabs | 1 | 623.08 |
| Fiber Optic Patch Cable | M107L02 | Thorlabs | 2 | 260.18 |
| RFP Excitation Filter | ET560/40x | Chroma Technologies | 1 | 365 |
| Excitation Filter Holder Part 1 | SM05SMA | Thorlabs | 1 | 35.59 |
| Excitation Filter Holder Part 2 | PM20-SMA | Thorlabs | 1 | 40.36 |
| LED Driver | LEDD1B | Thorlabs | 1 | 380.04 |
| LED Driver Power Cable | KPS201 | Thorlabs | 1 | 43.15 |
| Microcontroller for LED Control | FT232H | Adafruit | 1 | 14.95 |
| Digital to Analog Converter | BOB-12918 | Sparkfun | 1 | 6.95 |
| BNC Male Plug Terminal Block | 2888 | Adafruit | 1 | 0.95 |
| **One Color Blue** | | | | |
| DAPI LED | M375F3 | Thorlabs | 1 | 568.17 |
| Fiber Optic Patch Cable | M107L02 | Thorlabs | 2 | 260.18 |
| DAPI Excitation Filter | AT375/28x | Chroma Technologies | 1 | 365 |
| Excitation Filter Holder Part 1 | SM05SMA | Thorlabs | 1 | 35.59 |
| Excitation Filter Holder Part 2 | PM20-SMA | Thorlabs | 1 | 40.36 |
| LED Driver | LEDD1B | Thorlabs | 1 | 380.04 |
| LED Driver Power Cable | KPS201 | Thorlabs | 1 | 43.15 |
| Microcontroller for LED Control | FT232H | Adafruit | 1 | 14.95 |
| Digital to Analog Converter | BOB-12918 | Sparkfun | 1 | 6.95 |
| BNC Male Plug Terminal Block | 2888 | Adafruit | 1 | 0.95 |
| **2 Color** | | | | |
| Bifurcated Fiber Bundle | BFY1000LS02 | Thorlabs | 1 | 502.65 |
| Fiber Optic Patch Cable | M107L02 | Thorlabs | 2 | 260.18 |
| Excitation Filter Holder Part 1 | SM05SMA | Thorlabs | 2 | 35.59 |
| Excitation Filter Holder Part 2 | PM20-SMA | Thorlabs | 2 | 40.36 |
| LED Driver | LEDD1B | Thorlabs | 2 | 380.04 |
| LED Driver Power Cable | KPS201 | Thorlabs | 2 | 43.15 |
| Microcontroller for LED Control | FT232H | Adafruit | 1 | 14.95 |
| Digital to Analog Converter | BOB-12918 | Sparkfun | 2 | 6.95 |
| BNC Male Plug Terminal Block | 2888 | Adafruit | 2 | 0.95 |
| *Cost basis for all listed parts in in USD as of January 2026.* | | | | |

#### 2.6 LumeDEL LED and Fiber Optics Bill of Material 41

| *Identifier* | *Part Name/Number OR File Location* | *Source* | *Qty.* | *Cost Per* |
|---|---|---|---|---|
| **One Color Green** | | | | |
| 490nm Fiber Mounted LED | LumeDEL Model N490 (cat 70-903) | Edmund Optics | | |
| Fiber Optic Patch Cable | M107L02 | Thorlabs | 2 | 260.18 |
| GFP Excitation Filter | ET480/30x (12mm Circle Size) | Chroma Technologies | 1 | 365 |
| Excitation Filter Holder Part 1 | SM05SMA | Thorlabs | 1 | 35.59 |
| Excitation Filter Holder Part 2 | PM20-SMA | Thorlabs | 1 | 40.36 |
| **One Color Red** | | | | |
| 550nm Fiber Mounted LED | LumeDEL Model N550 (cat 70-905) | Edmund Optics | | |
| Fiber Optic Patch Cable | M107L02 | Thorlabs | 2 | 260.18 |
| RFP Excitation Filter | ET560/40x | Chroma Technologies | 1 | 365 |
| Excitation Filter Holder Part 1 | SM05SMA | Thorlabs | 1 | 35.59 |
| Excitation Filter Holder Part 2 | PM20-SMA | Thorlabs | 1 | 40.36 |
| **2+ Color** | | | | |
| Bifurcated Fiber Bundle | BFY1000LS02 | Thorlabs | 1 | 502.65 |
| Fiber Optic Patch Cable | M107L02 | Thorlabs | 2 | 260.18 |
| Excitation Filter Holder Part 1 | SM05SMA | Thorlabs | 2 | 35.59 |
| Excitation Filter Holder Part 2 | PM20-SMA | Thorlabs | 2 | 40.36 |
| LED Driver | LEDD1B | Thorlabs | 2 | 380.04 |
| LED Driver Power Cable | KPS201 | Thorlabs | 2 | 43.15 |
| Microcontroller for LED Control | FT232H | Adafruit | 1 | 14.95 |
| Digital to Analog Converter | BOB-12918 | Sparkfun | 2 | 6.95 |
| BNC Male Plug Terminal Block | 2888 | Adafruit | 2 | 0.95 |
| *Cost basis for all listed parts in in USD as of January 2026.* | | | | |

#### 2.7 Parts Tapping and Countersinking Guide

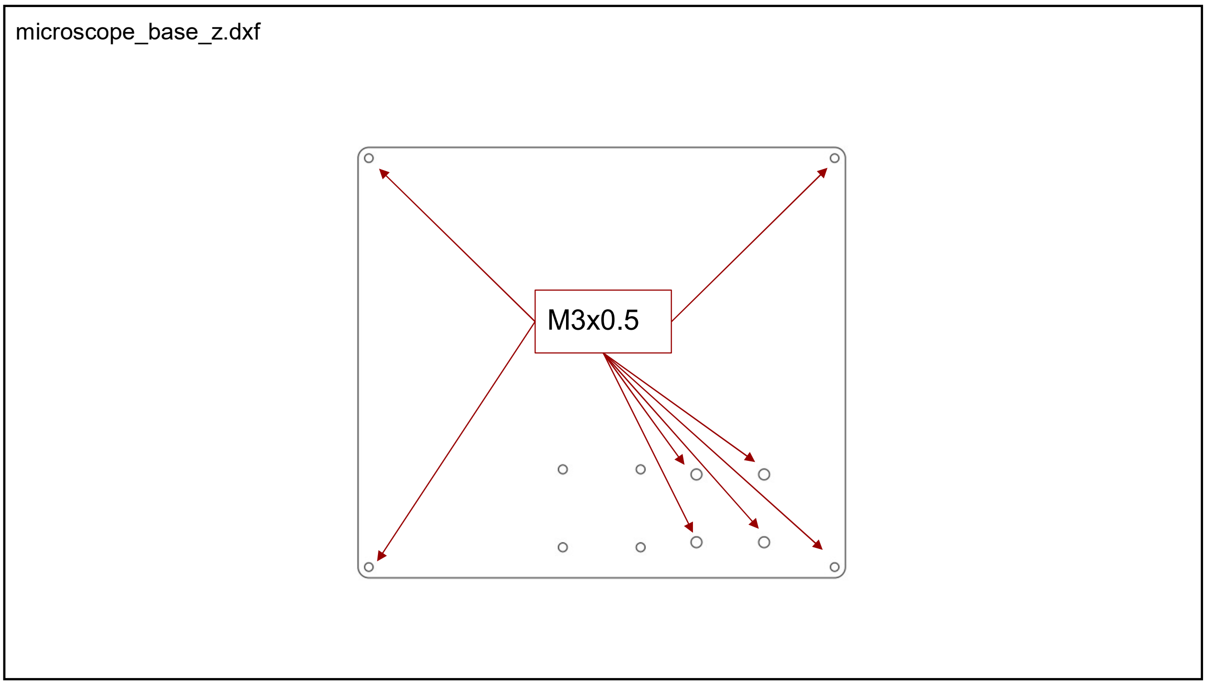

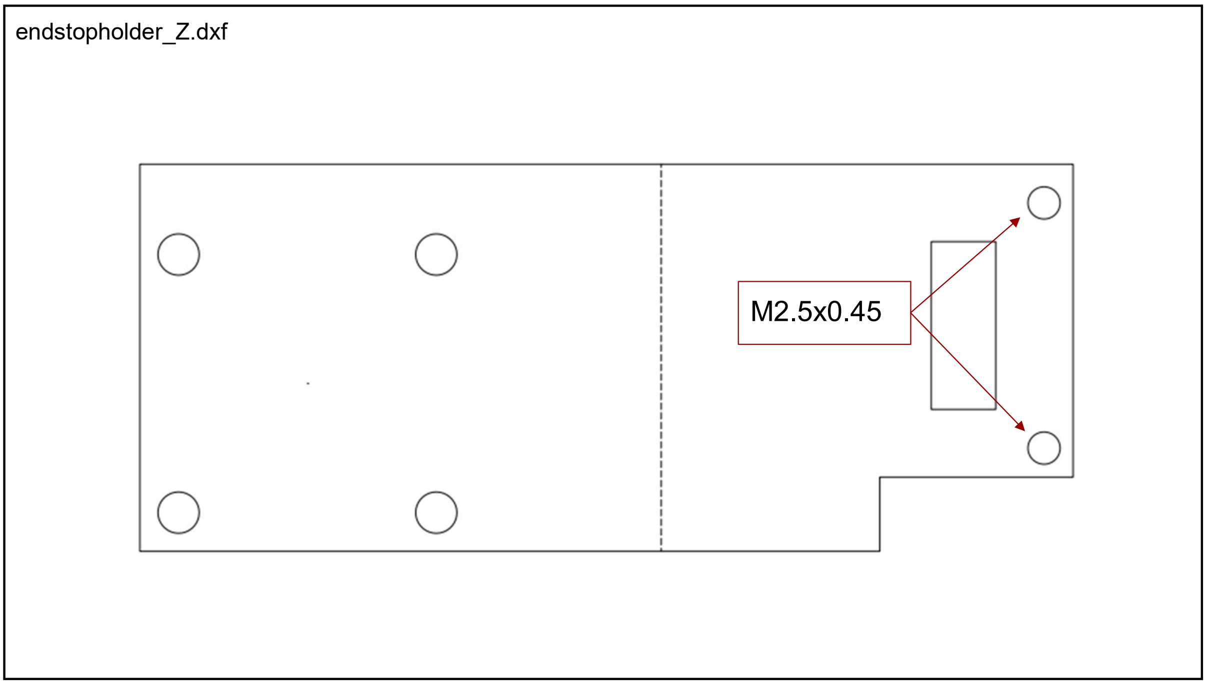

U_rail to alvium adapter.stp

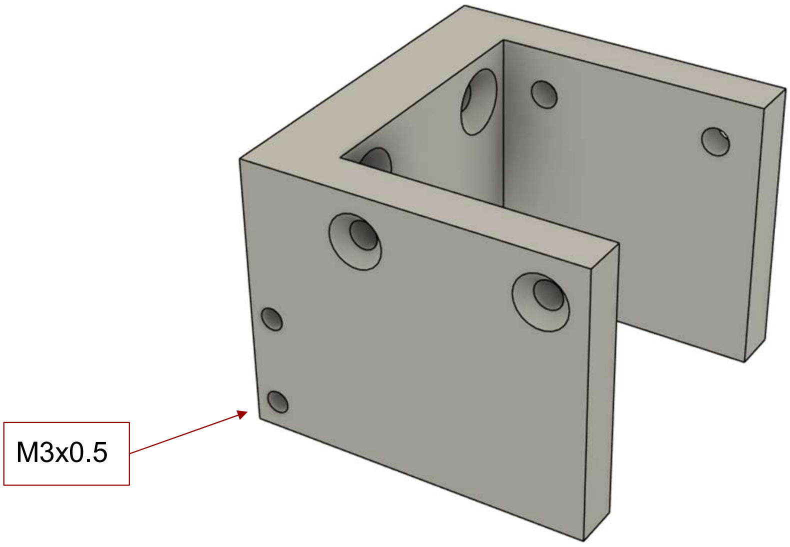

rotaryinner.stp

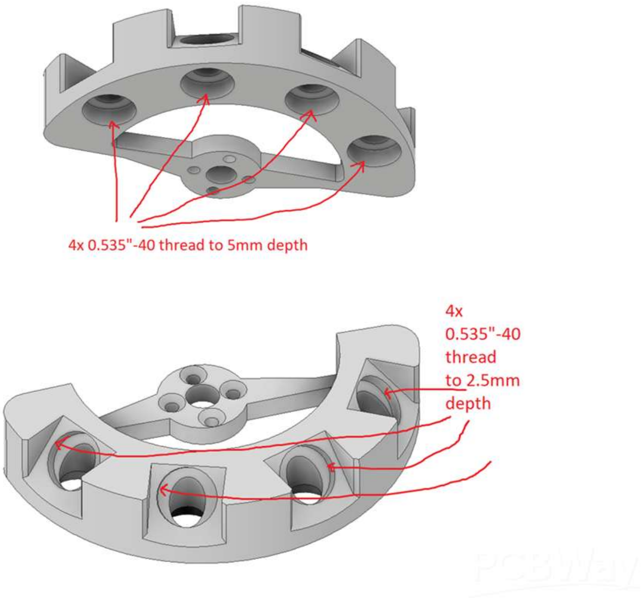

turret_outer_top.stp

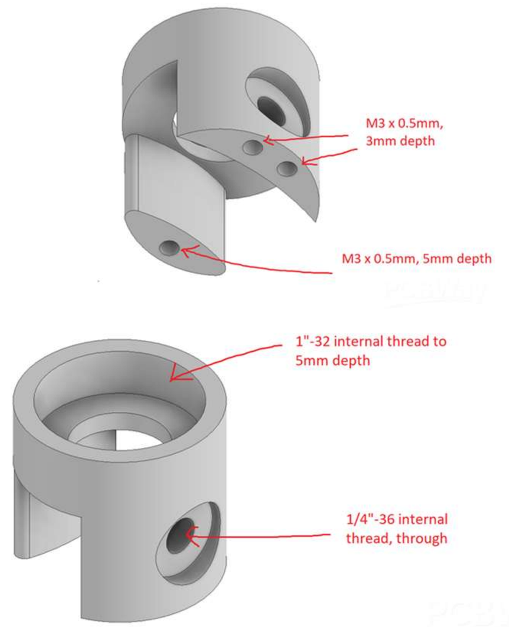

rotator to c-mount.stp

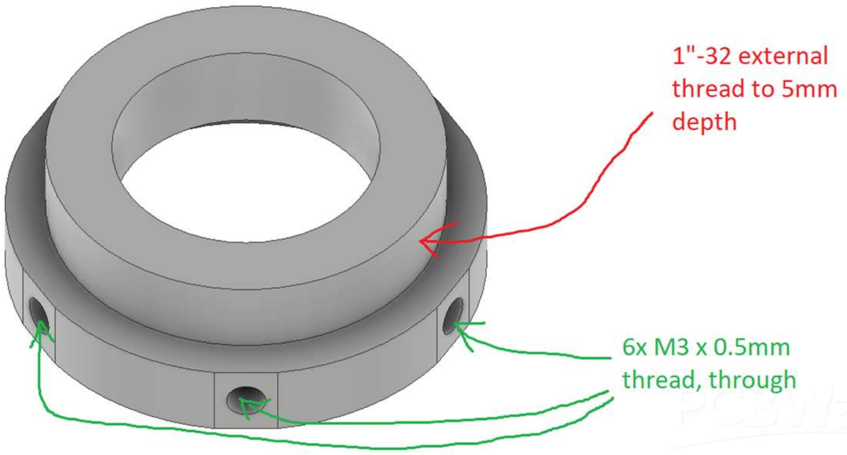

endstopholder_XYZ.stp

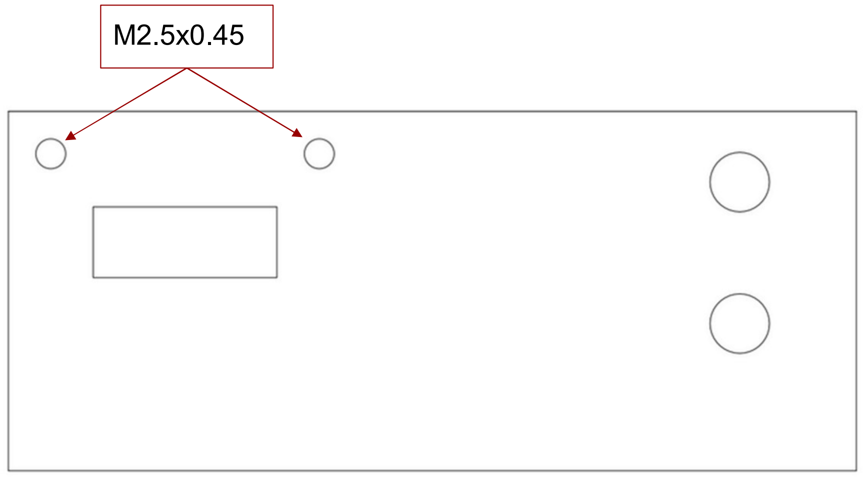

xyz_end_218_rev.dxf

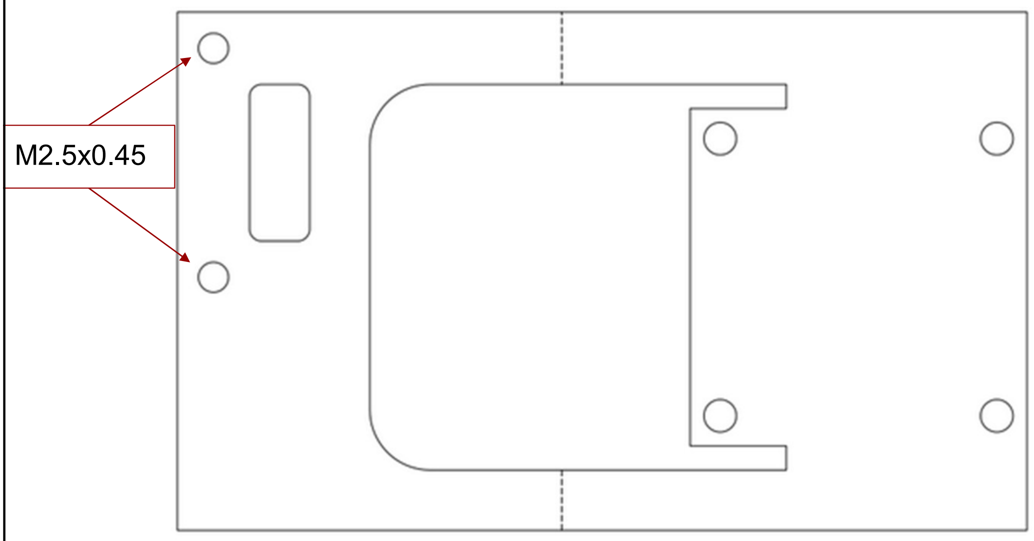

midbot_countersink_xyz.dxf

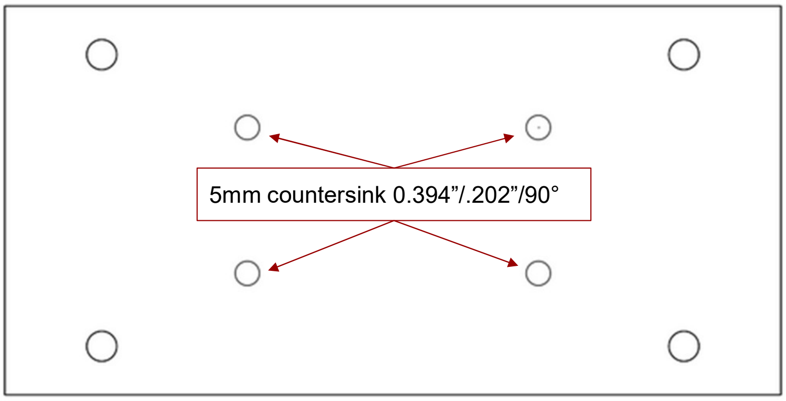

midbot_countersink_xyz.dxf

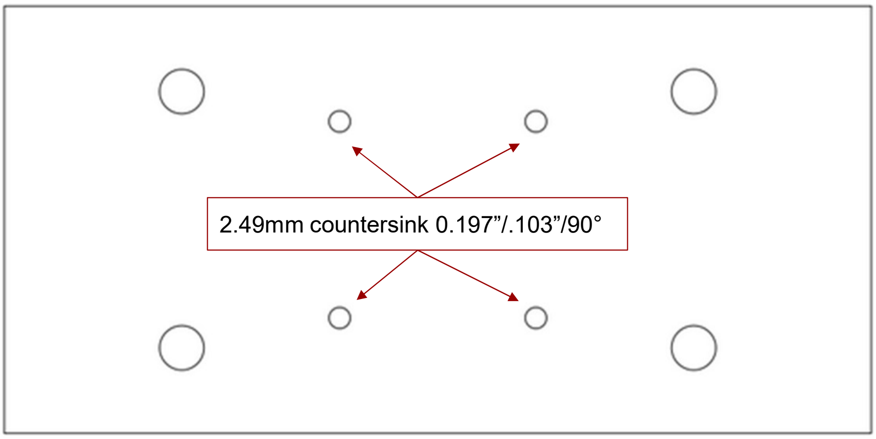

### 3 Supplementary Note 3: Assembly Guides

#### 3.1 Z Frame Build Guide

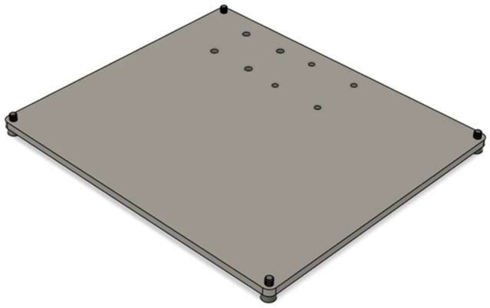

Thread in 4x M3x6 socket head screws into the corners of the base plate.

1

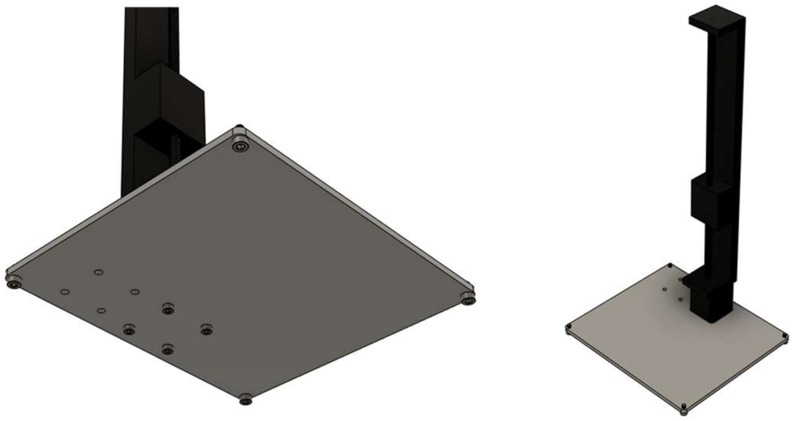

Remove the 4 M2.5 screws from the bottom of the linear rail. Using 4x M2.5x25 socket head screws, pin the base plate between the screw heads and the motor as shown in the right image.

2

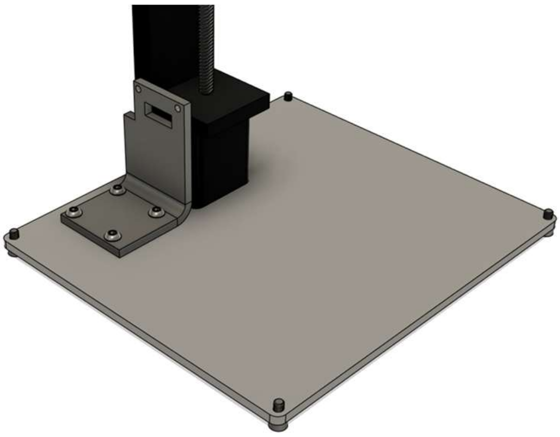

Mount the endstop holder using 4x M3x6 screws as shown in the picture.

3

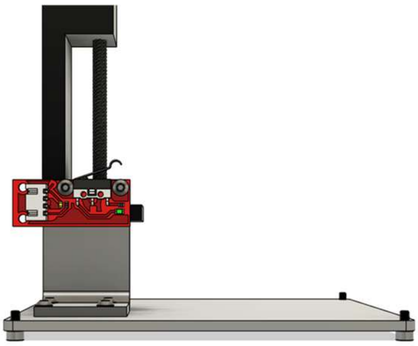

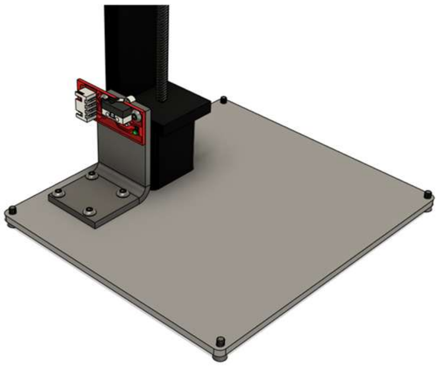

Mount the limit switch using 2x M2.5x4 screws as shown in the picture.

4

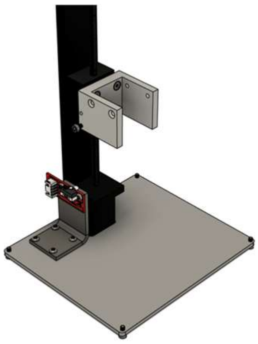 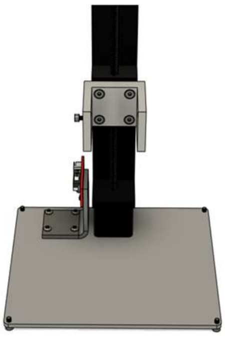

Mount the camera holder to the linear rail using 4x M3x6 flat head screws. Add the endstop hitting M3x8 socket head screw to the side of the camera holder as shown in the images.

5

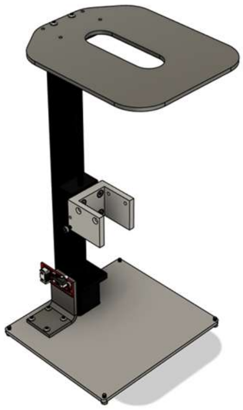

Remove the two M3 screws from the top of the linear rail and replace them with M3x10 screws, using them to secure the microscope sample plate in place.

6

#### 3.2 XYZ Frame Build Guide

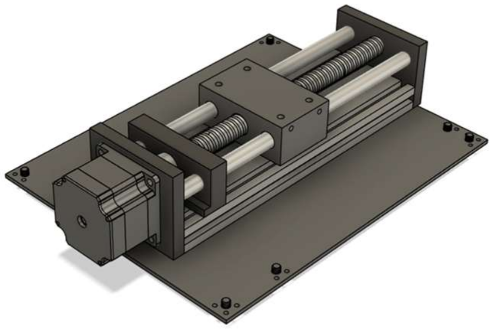

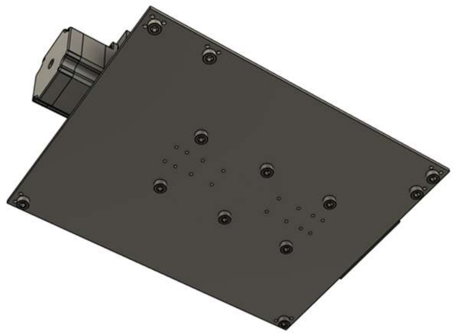

Secure the 150mm travel linear rail to the base using 6x M6x8 screws. Use the slot covers found in the bill of material to create spacers that align the sliding carriage nuts at the base of the motor to the holes on the sheet metal.

1

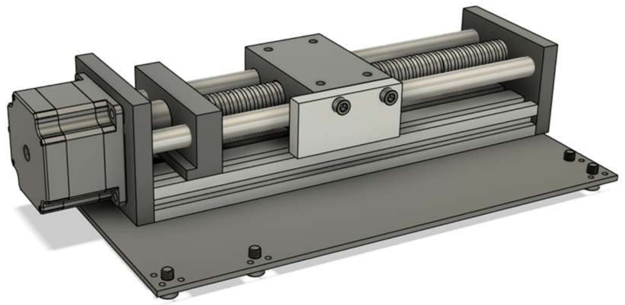

Mount the endstop hitter for the bottom axis in place using 2x M4x10 screws.

2

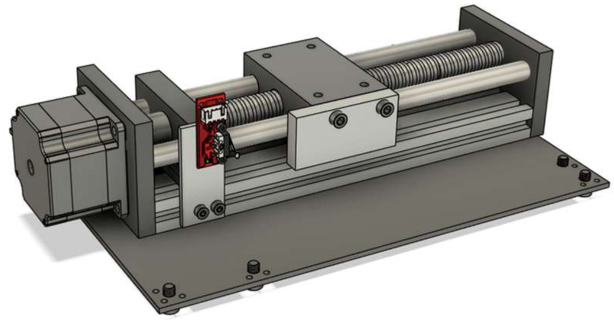

Mount the endstop assembly for the bottom axis in place using 2x M4x10 screws. To secure the limit switch, use 2x M2.5x4 screws.

3

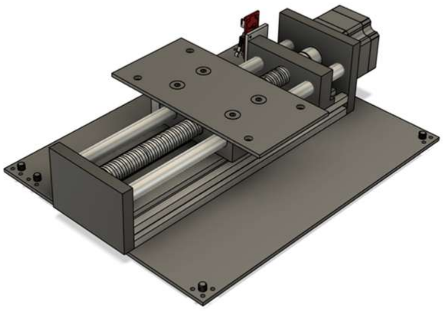

Mount the bottom to middle rail adapter plate in place using 4x M5x8 flathead screws.

4

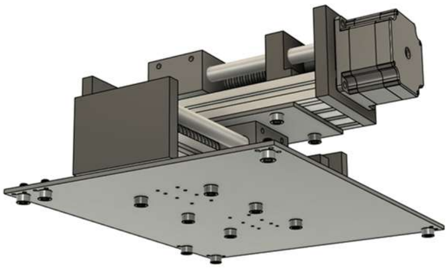 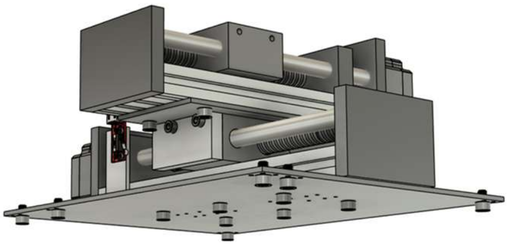

Secure the 100mm travel linear rail in place using 4x M6x8 screws. Use the slot covers found in the bill of material to create spacers that align the sliding carriage nuts at the base of the motor to the holes on the sheet metal.

5

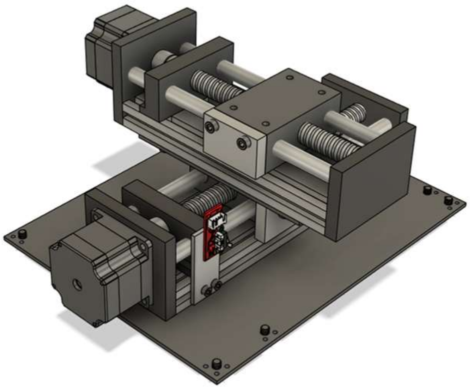

Mount the endstop hitter for the top axis in place using 2x M4x10 screws.

6

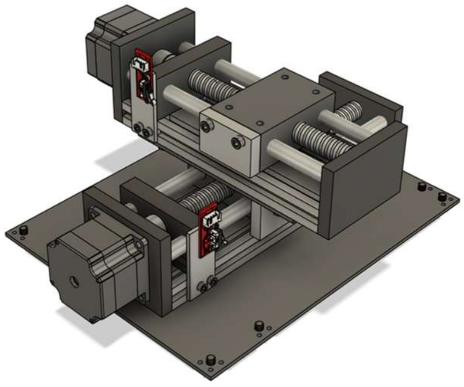

Mount the endstop assembly for the bottom axis in place using 2x M4x10 screws. To secure the limit switch, use 2x M2.5x4 screws.

7

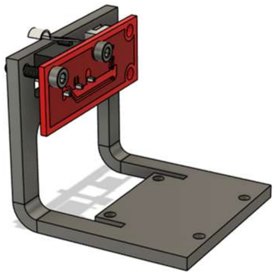

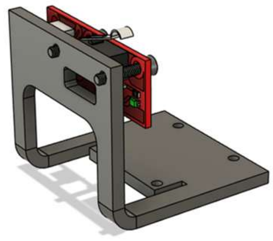

Secure the endstop to the bent endstop holder using 2x M2.5x10 screws.

8

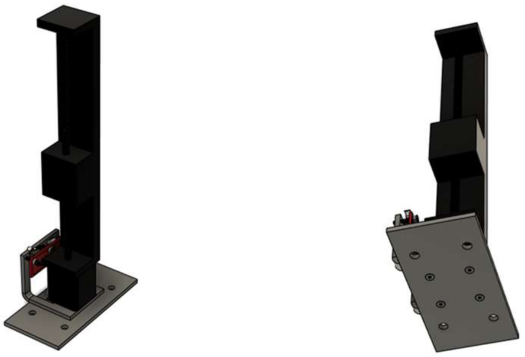

Secure the middle rail to top rail adapter to the base of the black linear rail using 4x M2.5x32 screws , pinning the endstop holder in place as shown in the images.

9

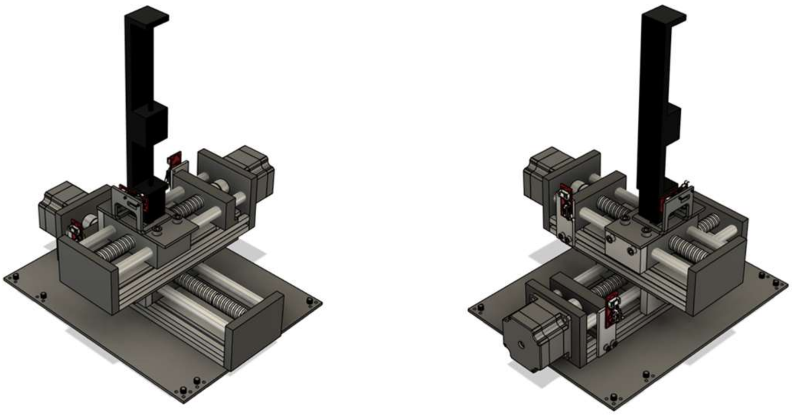

Secure the Z rail assembly from the previous steps to the middle linear rail using 4x M6x8 screws.

10

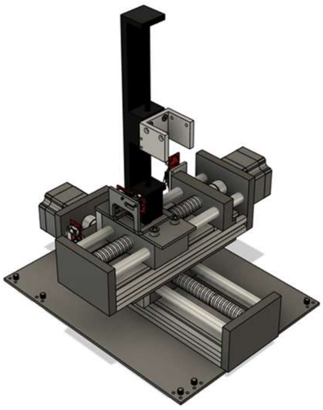 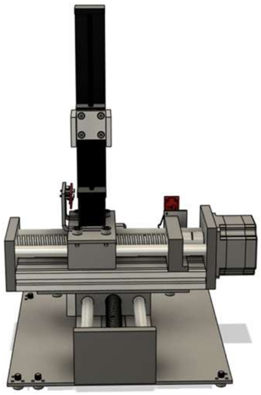

Mount the camera holder to the linear rail using 4x M3x6 flat head screws.

11

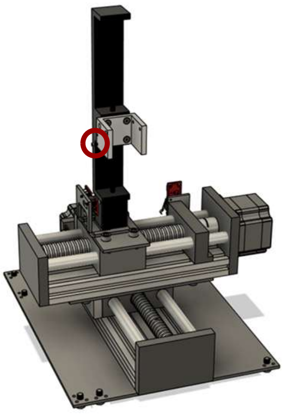

Add the endstop hitting M3x8 socket head screw to the side of the camera holder as shown in the images.

12

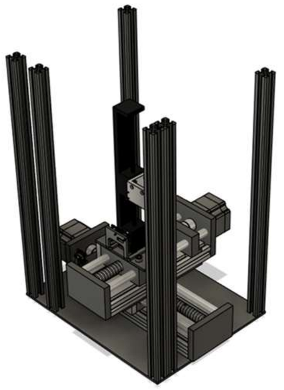

Secure the 400mm aluminum extrusions into place using 6 M6x8 screws as shown in the picture.

13

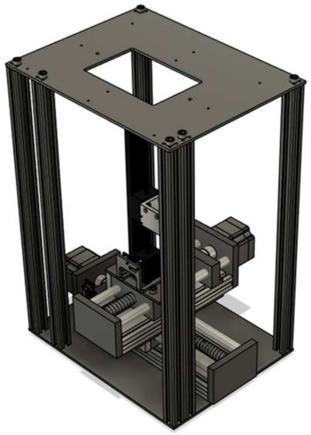

Secure lid of the system using 6x M6x8 screws as shown in the picture.

14

### 3.3 1 Color Optics Core Build Guide

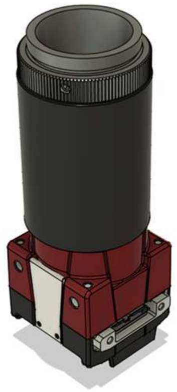

Mount the C-Mount lens barrel onto the camera, and then the rotating double sided C-Mount adapter on top of that. Complete this step with the camera pointing down to prevent particles from falling onto the sensor while you attach the parts.

1

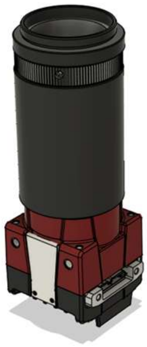

Screw on the SM1A10 C-Mount to SM1 Adapter to the top of the rotating double sided C-Mount adapter. Complete this step with the camera pointing down to prevent particles from falling onto the sensor while you attach the parts.

2

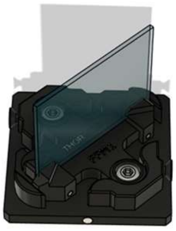

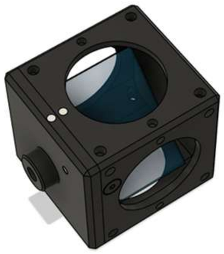

Open the filter cube and insert the dichroic mirror, taking note of the orientation marker supplied by the manufacturer on one of the sides of the mirror. Once assembly is done, make sure the marker is pointing the same direction as incoming excitatory light. The spring loaded jaws of the cube will hold the mirror in place. Reassemble the cube and use the screws you used to open it to close it.

3

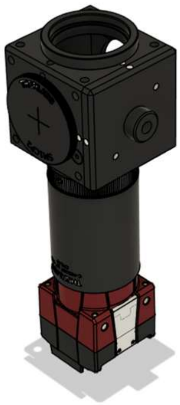

Mount the assembled filter cube on top of the C-Mount to SM1 adapter. Attach the SM1 cap to the face of the cube shown in the picture. Attach the SM1A9 SM1 to C-Mount adapter to the top of the cube

Use the set screw on the rotating double sided C-Mount adapter to loosen the rotating element and rotate it until it appears as show in the image.

4

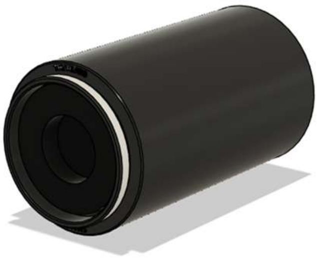

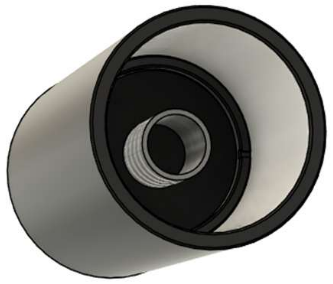

Take the SM1L20 lens barrel and thread a SM1 retaining ring all the way to the bottom. Then, add the collimator as shown in the left image. Finally, secure the assembly with another retaining ring as shown in the right image. Align the collimator such that the lens is in the middle of the barrel.

5

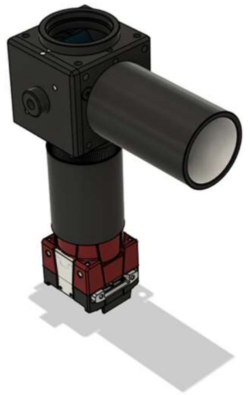

Mount the collimator assembly to the side of the cube shown in the picture. It should point directly at face of the dichroic mirror.

6

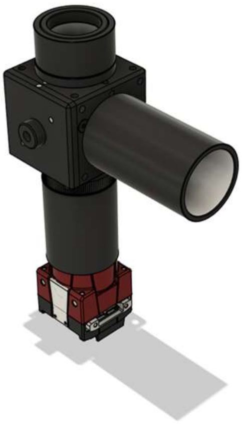

Mount the 15mm lens tube above the SM1 to C-Mount adapter, and then the C-Mount to RMS adapter to the top of that barrel.

7

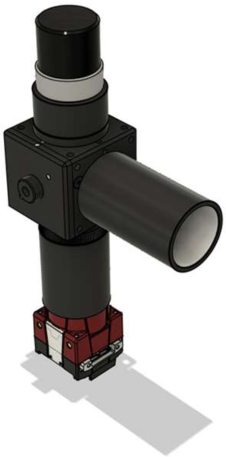

Add the objective to the top of the assembly.

8

#### 3.4 2 Color Optics Core Build Guide

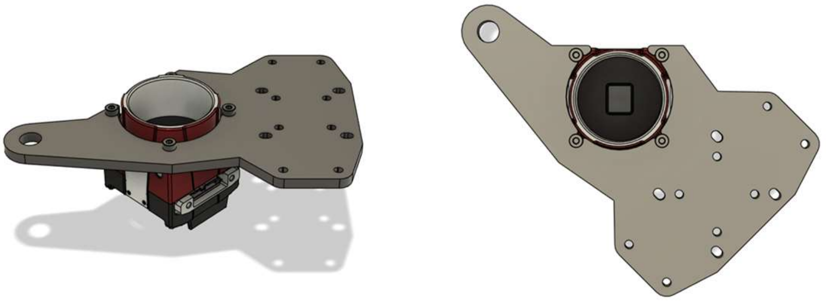

Mount the camera servo mount to the screw holes on the top of the camera in the orientation as shown in the images.

1

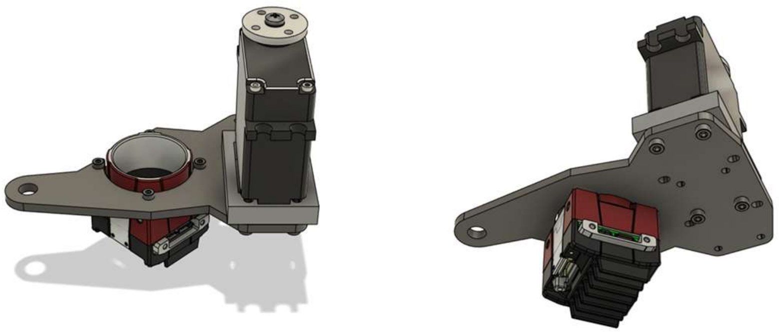

Mount the servo in place using the servo booster seat and 4x M2x16 screws in the arrangement shown in the pictures. Use the servo horn shown in the photo, it comes in the box with the servo.

2

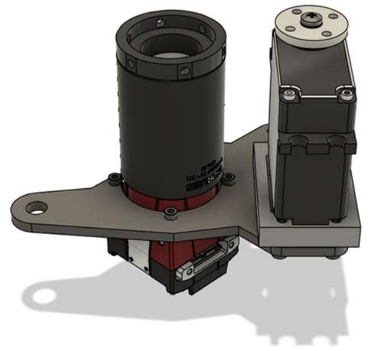

Mount the 40mm lens barrel to the camera, and the C-Mount to turner turret adapter above that.

3

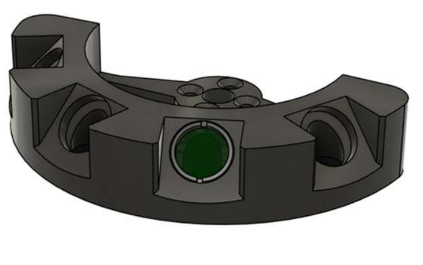

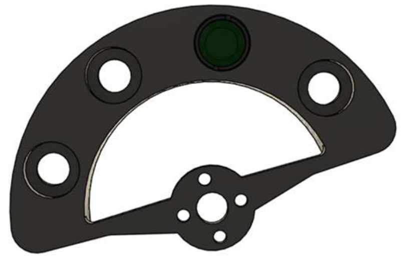

Use SM05 retaining rings to mount a dichroic mirror and an emission filter in place. Use the indicators on the side of the filter so show which way the filter should face, consider the direction light will travel. The dichroic mirror should be in the upper slot.

4

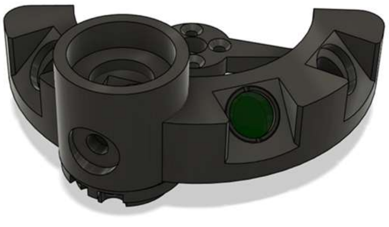

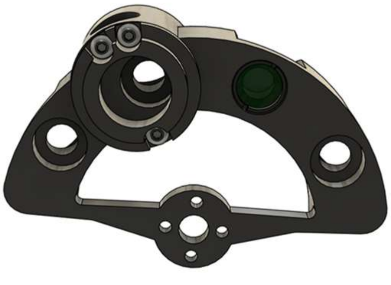

Mount the turret turner top and bottom in place using 3x M3x4 button cap screws as shown in the images.

5

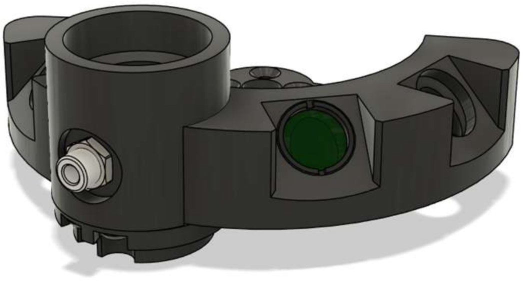

Add the HASMA SMA adapter to the front of the turner turret top, using the jam nut to secure it in place.

6

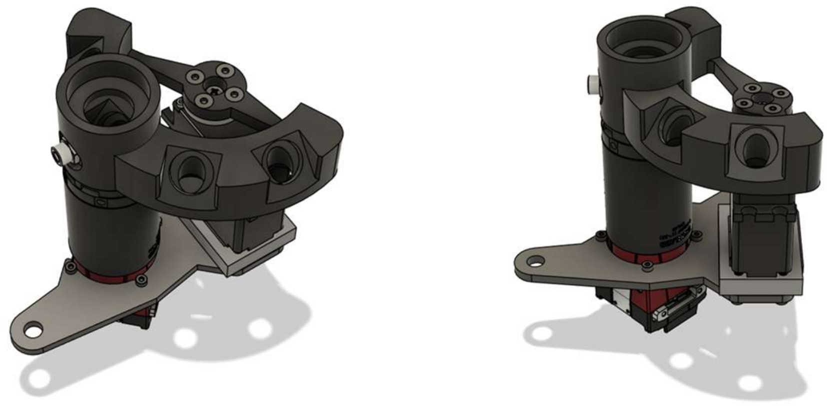

Align the turning assembly to the servo motor horn and 4x M3x6 flathead screws. The bottom of the turner assembly should sit flush with the part below it.

7

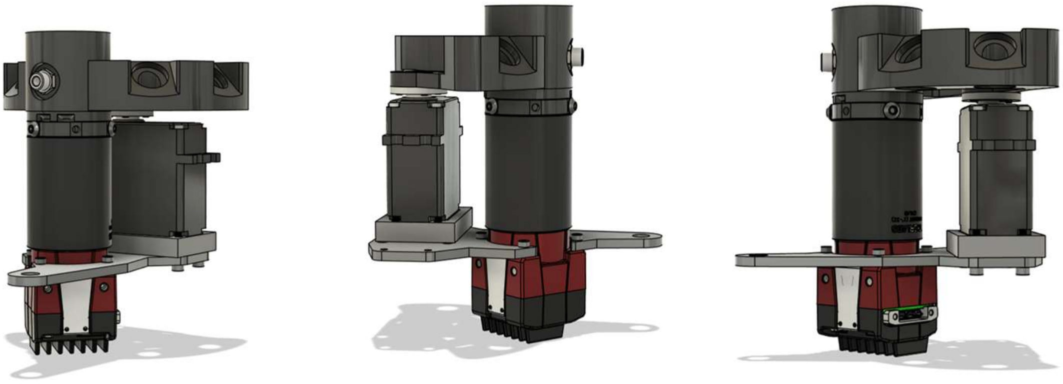

Secure the turning assembly in place using 3x M3x6 set screws on the sides of the turner assembly as shown in the pictures.

8

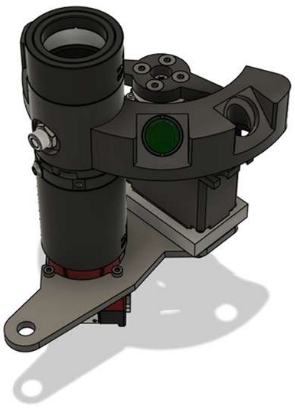

Add the 15mm C-Mount barrel above the turning assembly. Insert the C-Mount to RMS adapter to the top of that lens barrel as shown in the picture.

9

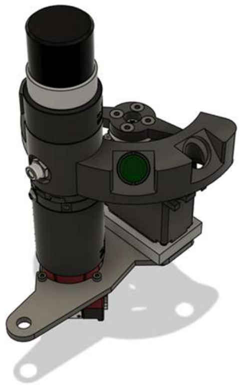

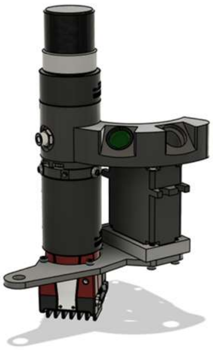

Add the objective to the top of the assembly.

10

#### 3.5 Module Changing Guide

48

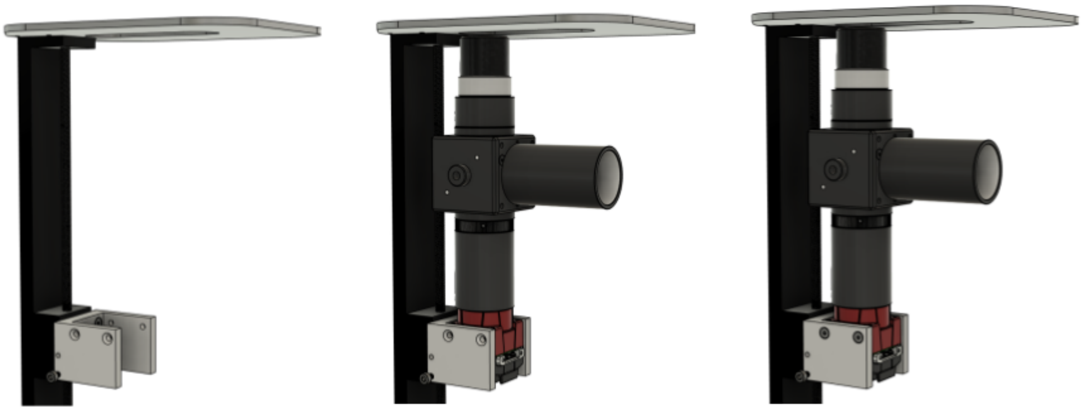

To swap out the optics module, slide in the camera assembly, line it up with the holes on the side of the camera adapter, and secure it in place with 4 M3x8 flathead screws.

### 4 Supplementary Note 4: Wiring Diagrams and Configuration

#### 4.1 1 Color Fiber Optic Cable Connection Diagram

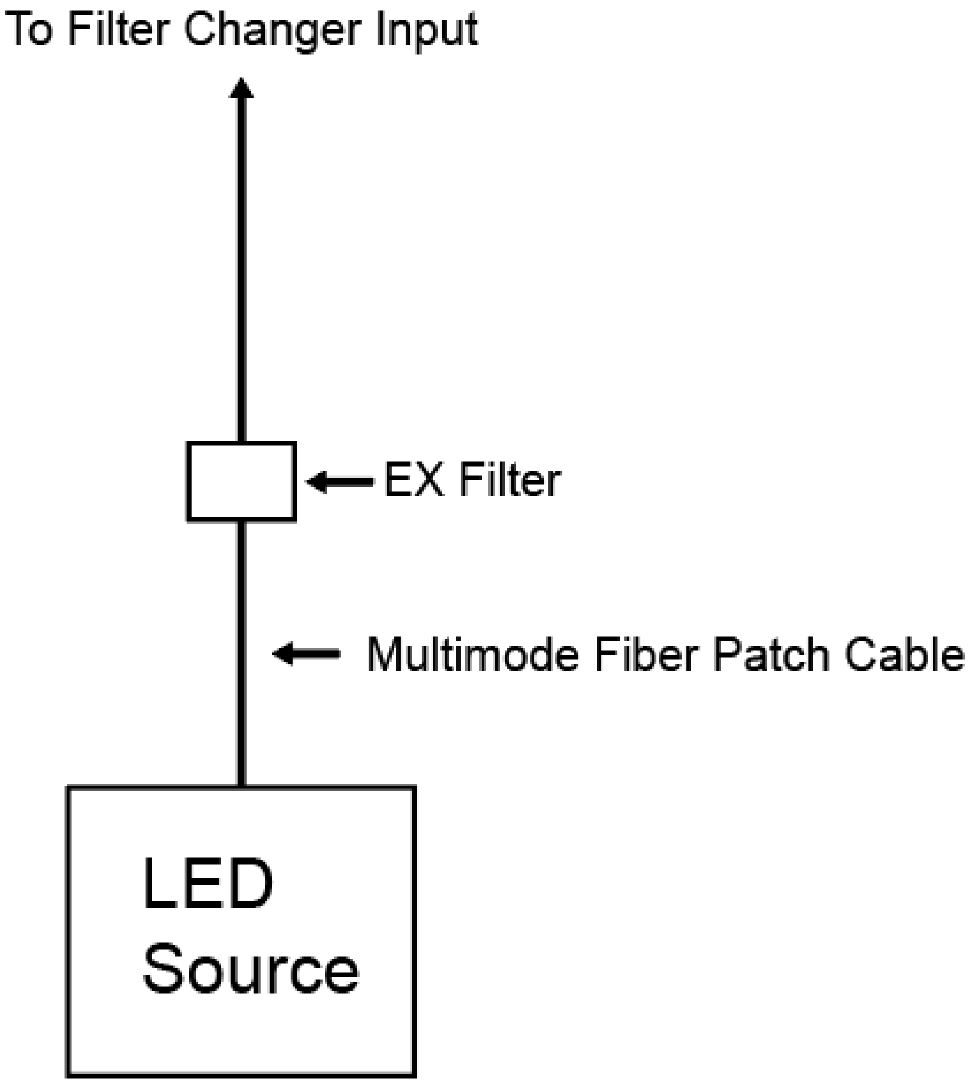

#### 4.2 2 Color Fiber Optic Cable Connection Diagram

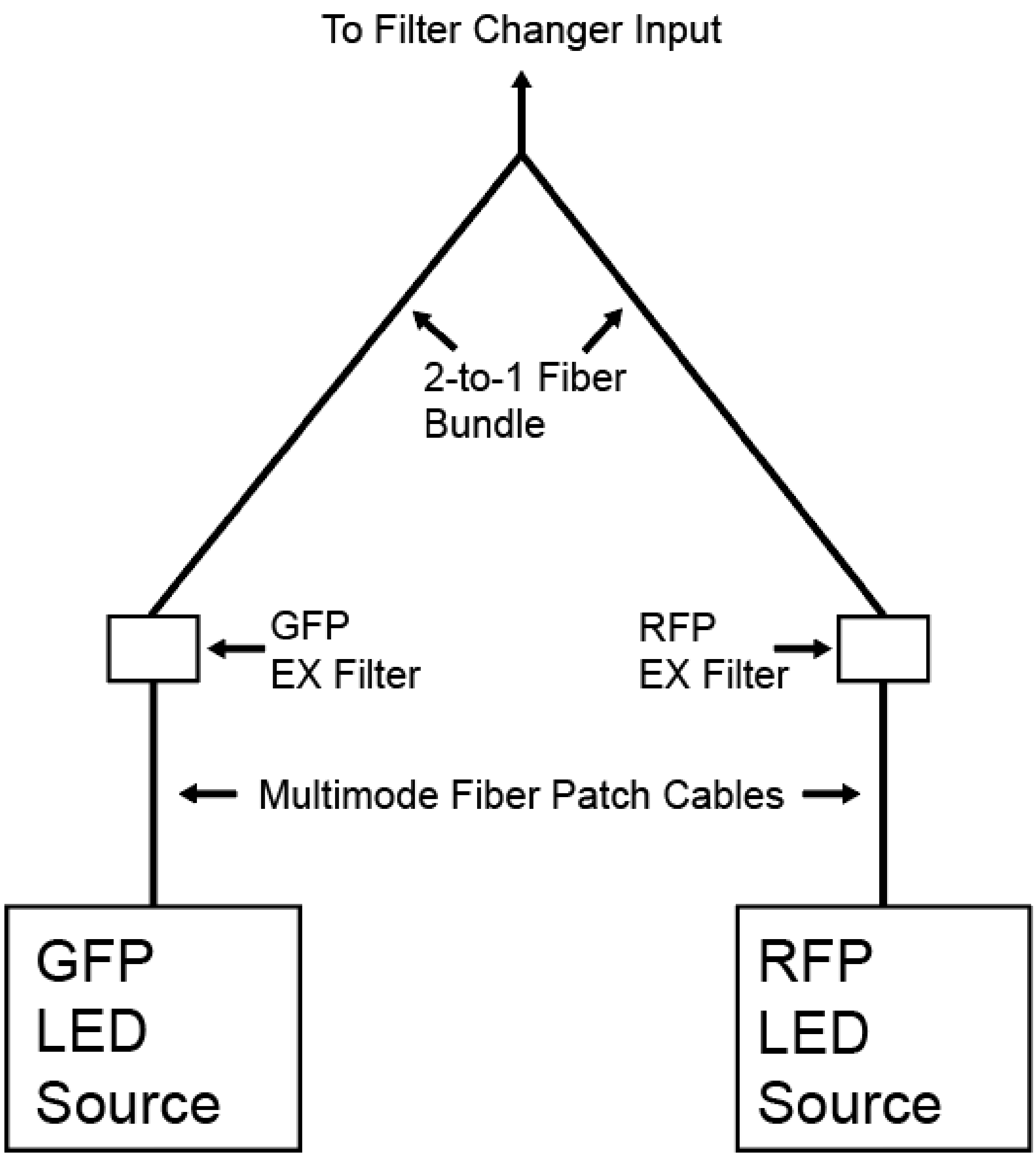

#### 4.3 1 Color Thorlabs Driver Wiring Diagram

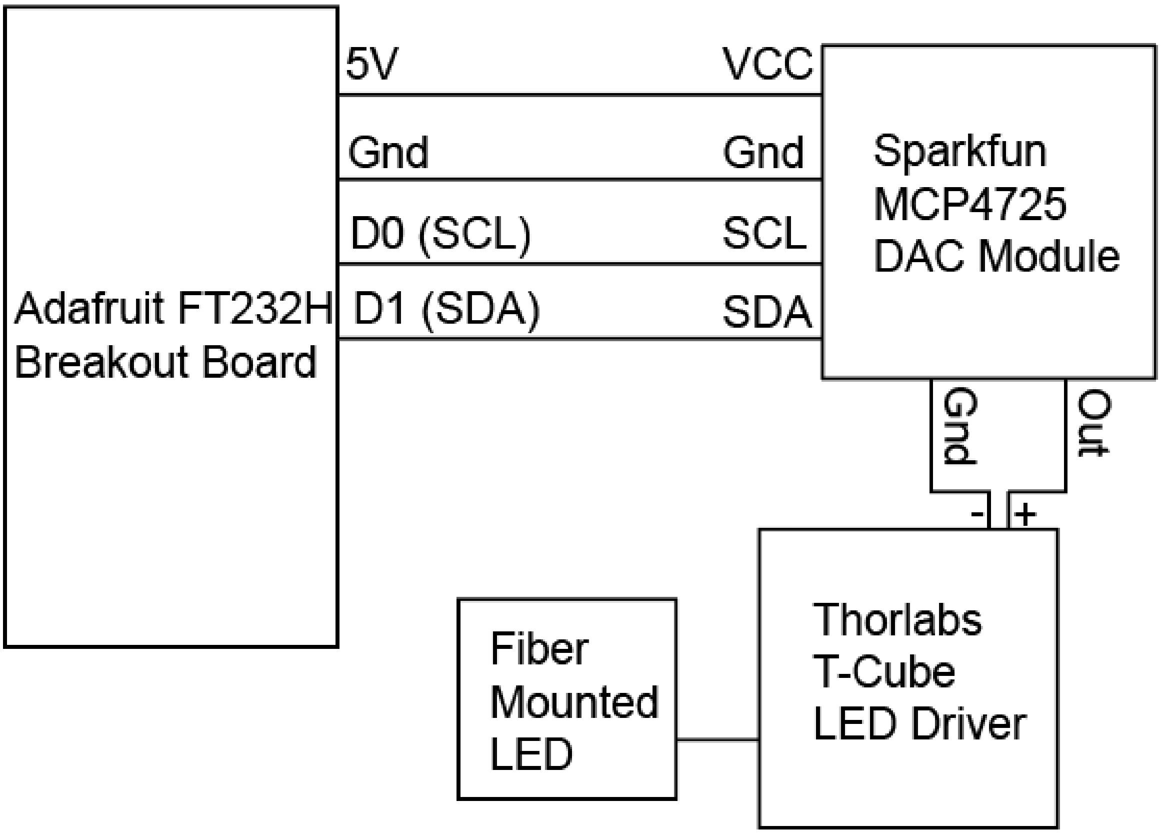

#### 4.4 2 Color Thorlabs Driver Wiring Diagram

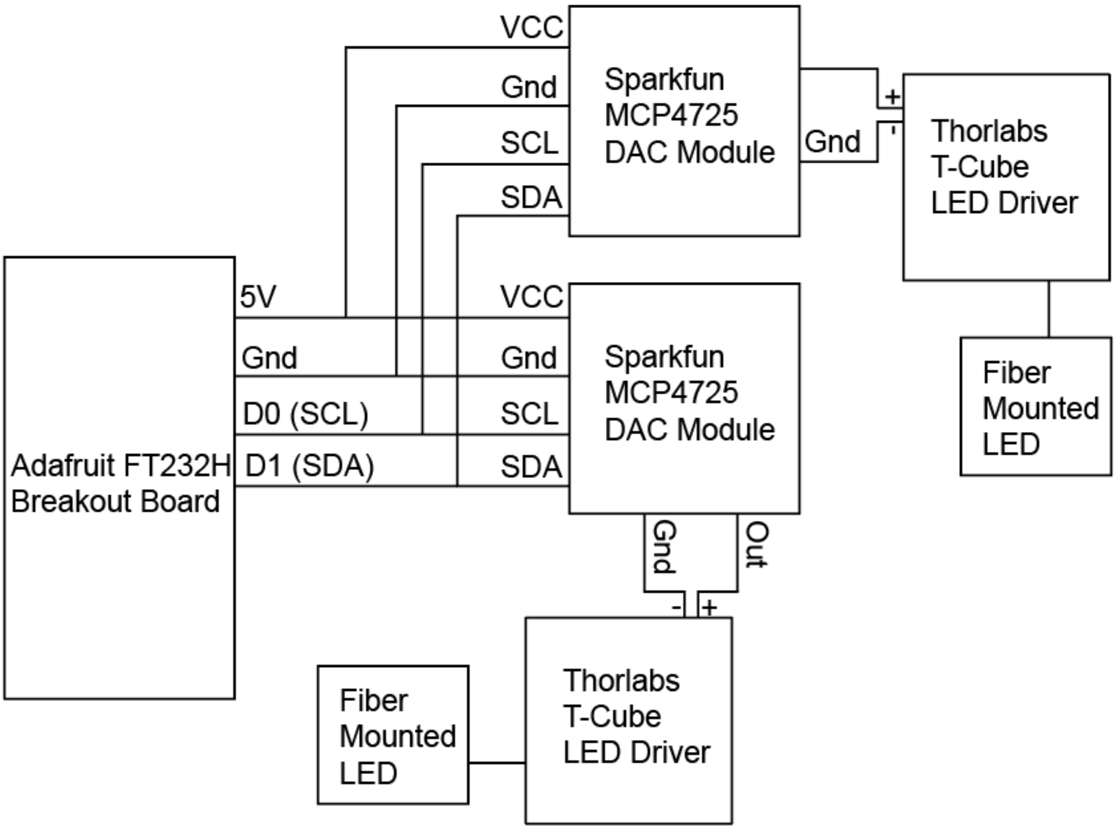

### 4.5 Polulu Tic T825 Stepper Driver Software Configuration 54

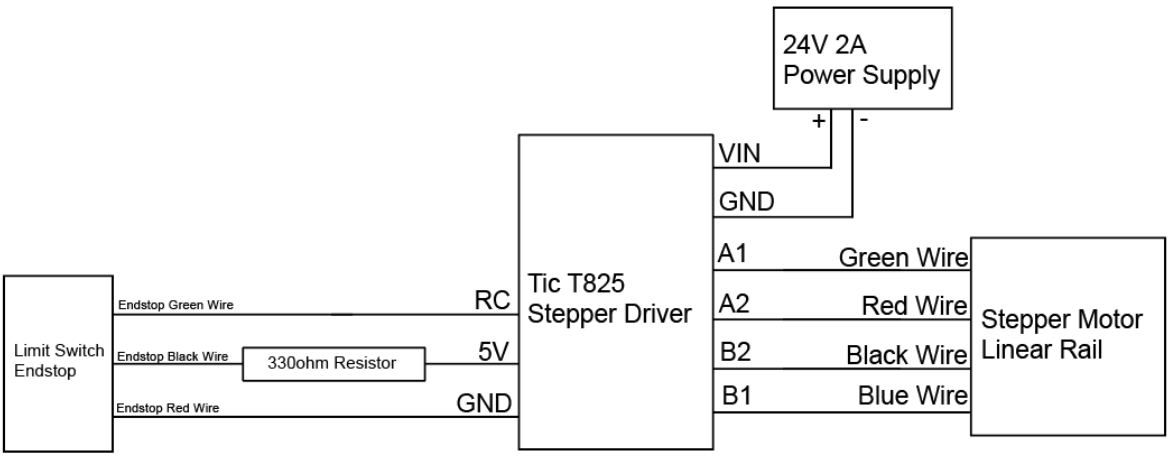

#### 4.6 Polulu Maestro Servo Driver Software Configuration

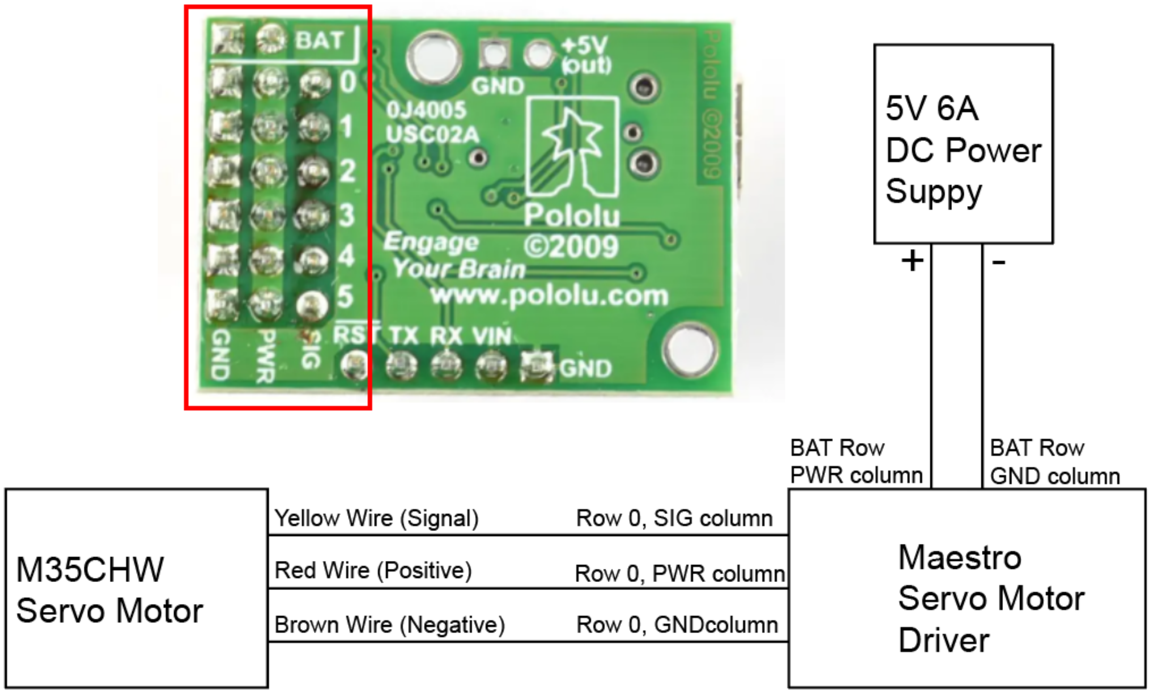

### 4.7 Z Axis Stepper Motor Configuration For All Systems 56

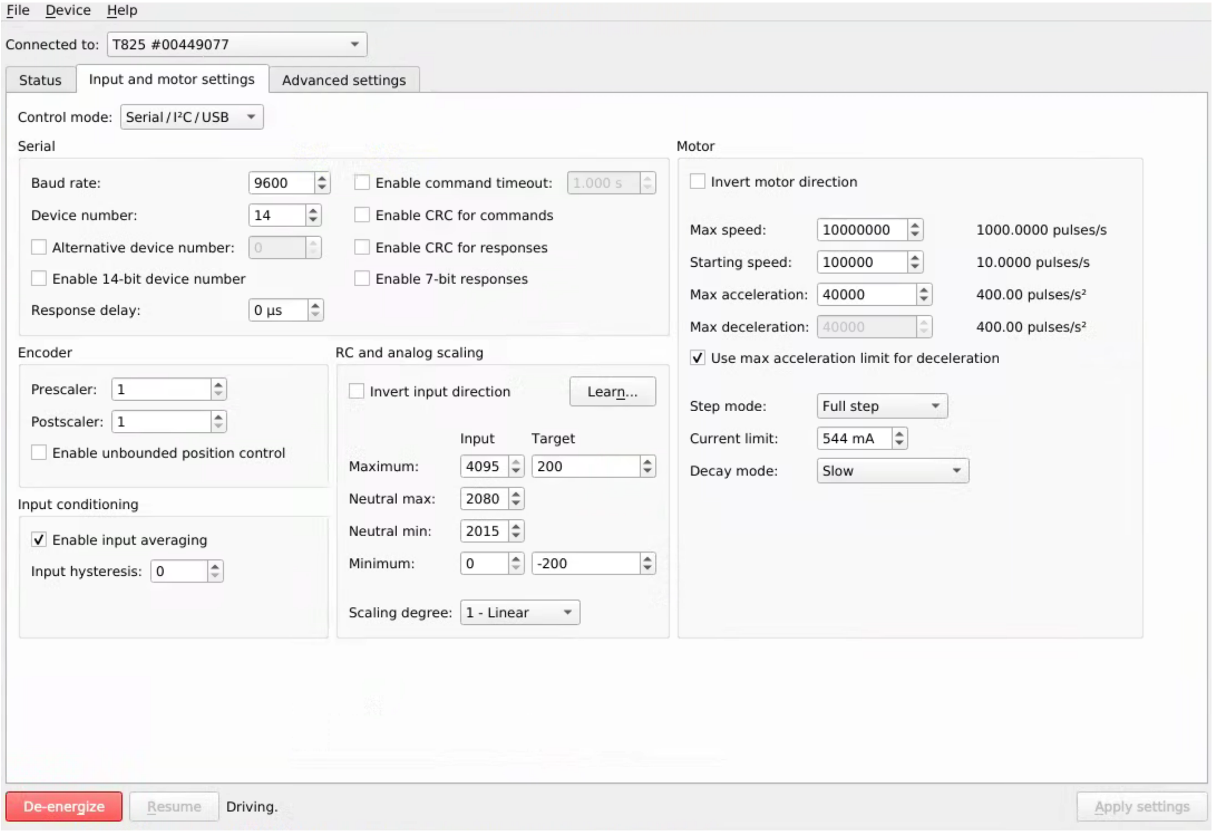

#### 4.8 X and Y Axis Stepper Motor Configuration For Three Axis Module 57

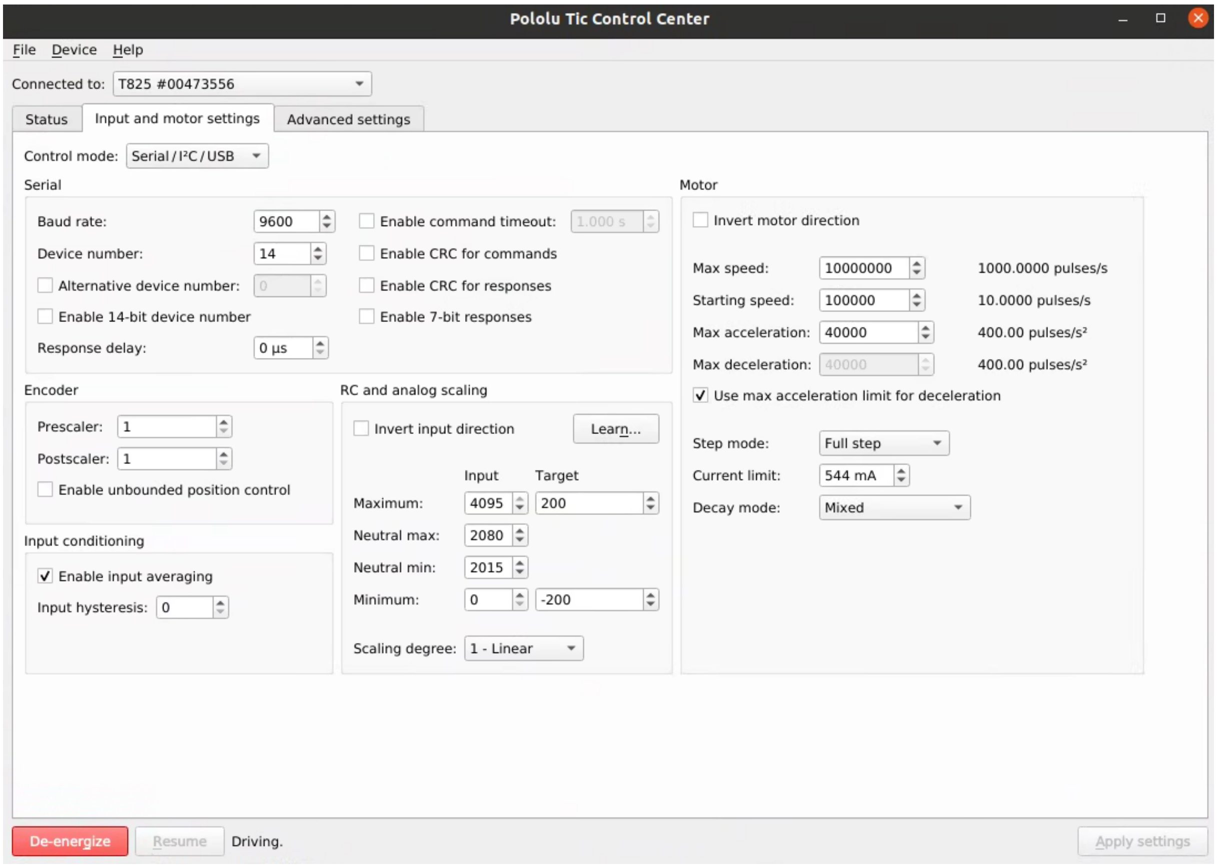

#### 4.9 Endstop Configuration For All Motors

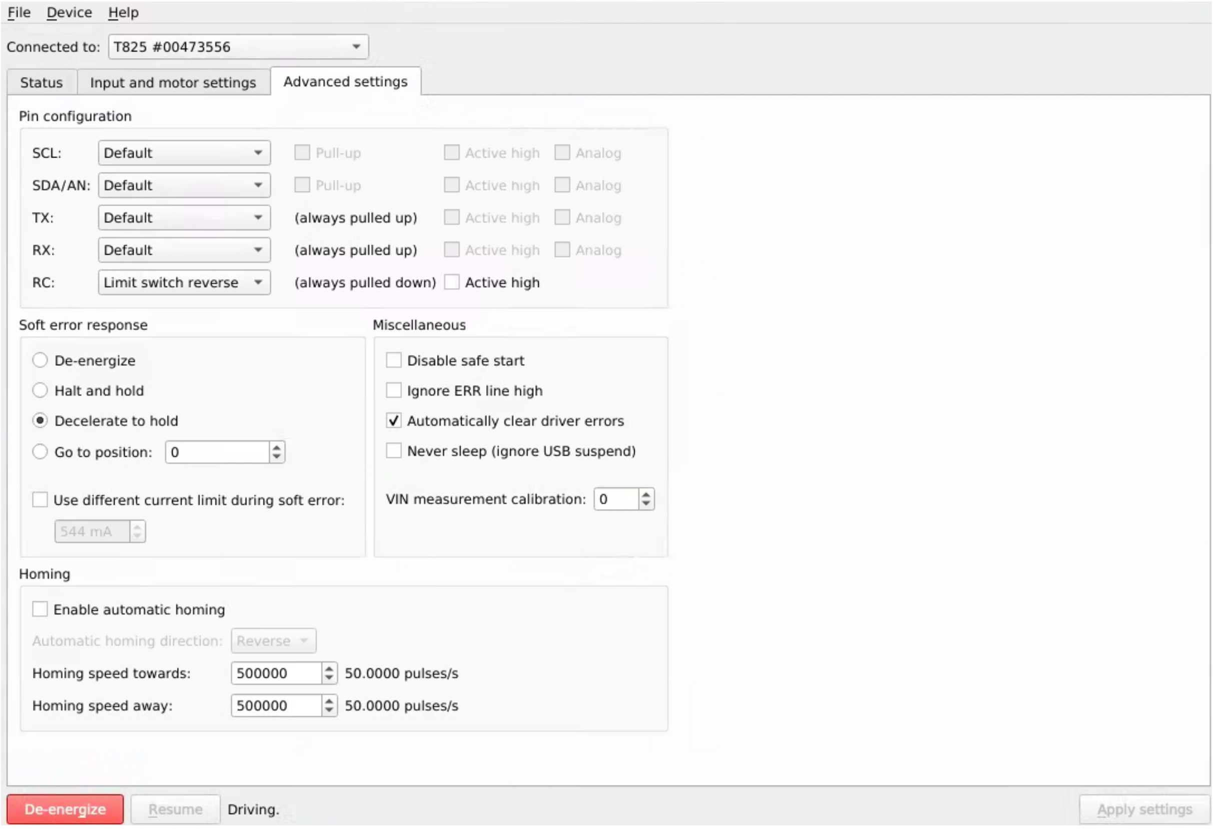

#### 4.10 Configuration For Filter Turret Servo Motor

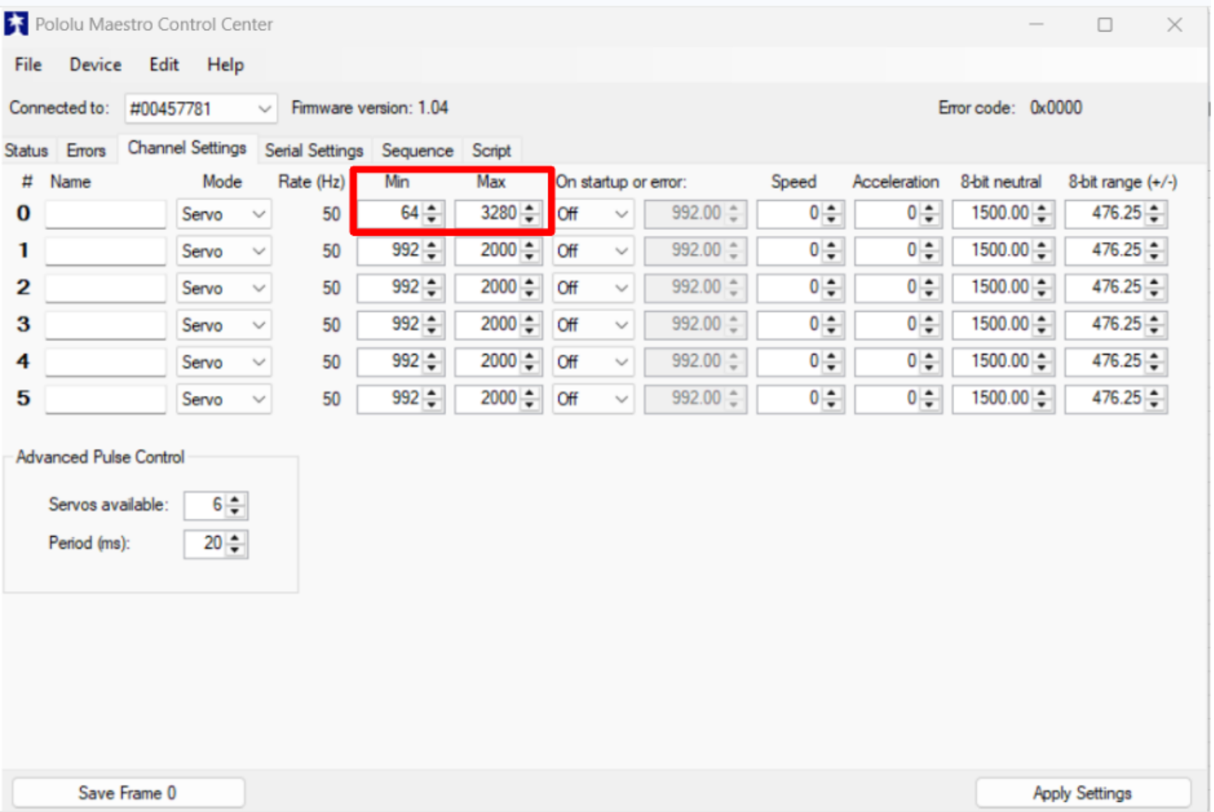
